# The first fossil skull of an anteater (Vermilingua, Myrmecophagidae) from northern South America, a taxonomic reassessment of *Neotamandua* and a discussion of the myrmecophagid diversification

**DOI:** 10.1101/793307

**Authors:** Kevin Jiménez-Lara, Jhon González

## Abstract

The evolutionary history of the South American anteaters, Vermilingua, is incompletely known as consequence of the fragmentary and geographically biased nature of the fossil record of this group. *Neotamandua borealis* is the only recorded extinct species from northern South America, specifically from the Middle Miocene of La Venta area, southwestern Colombia. A new genus and species of myrmecophagid for La Venta, Gen. et sp. nov., is here described based on a new partial skull. Additionally, given that the co-occurrent species of Gen. et sp. nov., *N. borealis*, was originally referred to as *Neotamandua*, the taxonomic status of this genus is revised. The morphological and taxonomic analyses of these taxa indicate that Gen. et sp. nov. may be related to *Tamandua* and that the justification of the generic assignments of the species referred to as *Neotamandua* is weak or insufficient. Two species previously referred to as *Neotamandua* (*N. magna* and *N.*? *australis*) were designated as *species inquirendae* and new diagnostic information for the redefined genus and its type species, *N. conspicua*, is provided. Together, these results suggest that the diversification of Myrmecophagidae was taxonomically and biogeographically more complex than what has been proposed so far. Considering the new evidence, it is proposed a synthetic model on the diversification of these xenartrans during the late Cenozoic based on the probable relationships between their intrinsic ecological constraints and some major abiotic changes in the Americas.

## Introduction

The anteaters of the suborder Vermilingua are part of Xenarthra, one of the more inclusive clades in the evolutionary tree of the placental mammals (Eutheria) and a characteristic group in the land mammal assemblages of the middle-late Cenozoic of the Americas (McDonald et al., 2008; Foley et al., 2016; Halliday et al., 2016; Feijoo and Parada, 2017). Within Xenarthra, Vermilingua belongs to Pilosa, a clade that also includes the sloths, i.e., Tardigrada. Today, Vemilingua comprises the genera *Cyclopes* (pygmy anteaters), *Tamandua* (collared anteaters) and *Myrmecophaga* (giant anteaters). These genera groups ten extant species, most of which (seven) belong to *Cyclopes*, according to the most recent exhaustive taxonomic revision (Miranda et al., 2017). The classic phylogenetic hypothesis reunites *Tamandua* and *Myrmecophaga* in the family Myrmecophagidae, while *Cyclopes* is located in a basal position with respect to Myrmecophagidae as the only recent form of the family Cyclopedidae (Engelmann, 1985). With the connotation of a superior taxonomic hierarchy (i.e., at the family level; Barros et al., 2008; Gibb et al., 2015), according to an early evolutionary divergence (Hirschfeld, 1976; Delsuc et al., 2001; Gibb et al., 2016) and in recognition of a more extended use in the scientific literature, the names Myrmecophagidae and Cyclopedidae are used here, instead of Myrmecophaginae and Cyclopinae *sensu* Gaudin and Branham (1998), respectively. However, the taxonomic content of Myrmecophaginae and Cyclopinae, including extinct forms, is considered as transferable to their counterparties (McDonald et al., 2008).

The living anteaters, whose mean body mass ranges from ∼0.4 to 30 kilograms (Gaudin et al., 2018), are highly, morphologically specialized mammals due to their remarkable skeleton and soft-anatomy modifications, which are closely related to their myrmecophagous diets, i.e., diets consisting of at least 90% of ants/termites (Redford, 1987; McDonald et al., 2008). Many of these adaptations, anatomically located in the skull and jaws, are associated between themselves in several ways, being part of the architecture of an integrated functional system of food apprehension and ingestion. Among these features, the following are some of the most noteworthy: rostral elongation and narrowing, basicranial-basifacial axis curvature, complete loss of teeth, gracile jaw, reduction of the adductor jaw muscles, unfused jaw symphysis and protrusible long tongue (Reiss, 2001; Gaudin and McDonald, 2008; McDonald et al., 2008). Several of these morphological specializations are convergent with those described for other myrmecophagous mammals such as the pangolins (Pholidota) and the aardvarks (Tubulidentata), so it is not surprising that early systematic researchers erroneously proposed close common ancestry of Vermilingua with these Old world groups based on their superficial similarities (e.g., Engelmann, 1978; Norman and Ashley, 1994).

In spite of their unique biology and ecology, at least in the context of the land mammals of the Americas, the evolutionary history of the anteaters is largely obscured by their poor, fragmentary and geographically biased fossil record (Hirschfeld, 1976; Gaudin and Branham, 1998; McDonald et al., 2008). Generally, five valid genera and nine species are recognized in the fossil record of Vermilingua, two genera and two species of which have extant representatives, i.e., *Myrmecophaga tridactyla* and *Tamandua tetradactyla*. Myrmecophagidae groups nearly all of these fossil taxa (only one genus and one species for Cyclopedidae) in a general biochron that begin ca. 18 million years ago (Mya), most of them distributed throughout the Neogene (McDonald et al., 2008). But while the record of this family for the latter period is taxonomically more diverse than that for the Quaternary, it also poses more difficulties in the systematic framework of the implicated taxa. The oldest member of Myrmecophagidae is *Protamandua rothi*, from the late Early Miocene of the Province of Santa Cruz, southern Argentina (Ameghino, 1904). This species has been well validated from a pair of incomplete skulls and several postcranial bones, but the validity of other co-occurrent putative vermilinguan (myrmecophagid?) taxa is, at least, questionable (Hirschfeld, 1976; McDonald et al., 2008). For the early Middle Miocene has been reported a myrmecophagid doubtfully assigned to *Neotamandua*, and yet used to create a new species from isolated humeral remains (*N.*? *australis*; Scillato-Yané and Carlini, 1998). In the latter genus, postcranial material of a medium-to-large sized anteater from La Venta area, southwestern Colombia, was also allocated with some uncertainty (Hirschfeld, 1985). The description of this material represents the only nominal extinct species from northern South America. *Neotamandua* chronologically extends to the Late Miocene and Early Pliocene with the species *N. magna* (Ameghino, 1919), *N. greslebini* (Kraglievich, 1940) and *N. conspicua* (type species; Rovereto, 1914), all of which come from northwestern Argentina (Provinces of Catamarca and Tucumán). This genus is typically recognized as morphologically similar (even directly ancestral) to *Myrmecophaga*, although the latter is smaller in body size (Hirschfeld, 1976; Gaudin and Branham, 1998). Considering the very few anatomically correlatable elements on which the different species referred to as *Neotamandua* are based, Hirschfeld (1976) and Scillato-Yané and Carlini (1998) suggested that this genus could be paraphyletic. Furthermore, the latter authors proposed the hypothesis that *Neotamandua* is composed by two distinct evolutionary lineages: the one more closely related to *Myrmecophaga* and other to *Tamandua*. In turn, these two lineages would have diverged in allopatry in South America, in such a way that the geographic origin of *Myrmecophaga* is located in northern South America, while that of *Tamandua* is in southern South America.

In this article, we describe the first fossil skull of a myrmecophagid (and vermilinguan) from northern South America. This specimen was collected in the Middle Miocene of La Victoria Formation of La Venta area, Colombia. This allowed us to revise the taxonomic status of *Neotamandua*, as it is the only nominal taxon previously reported for the same region and geological unit in Colombia. The results prompt the development of a discussion on a model of diversification for Myrmecophagidae in which new and previous hypothesis about this evolutionary event are synthesized. This contribution is intended to revaluate, expand and integrate biotic and abiotic evidence related to the diversification of this fascinating mammal group, emphasizing on the biogeographic role of tropical, low latitude regions of the Americas.

## Material and methods

The cranial specimen described for the first time here for Colombia (VPPLT 975) comes from a light-brown mudstone layer in the Llano Largo field, around 2 Km northeast of La Victoria town, Municipality of Villavieja, Department of Huila (Fig. 1). Strata of the La Victoria Formation outcrop there, within the paleontologically relevant area of La Venta. La Victoria Formation is a geological unit of ∼500 m in thickness which is mainly composed by bioturbated mudstones (Anderson et al., 2016). These sedimentites are interrupted by very continuous, coarse-to-fine grained sandstones with crossbedding and erosive bases. According to the lithostratigraphic scheme of Guerrero (1997; Fig. 1), the new skull was found at a level stratigraphically close (<20 m) and below the Chunchullo sandstone beds, i.e., the lower part of the La Victoria Formation. This corresponds to the unit referred to as “Unit between the Cerro Gordo and Chunchullo sandstone beds”. As described by the same author, this unit, whose thickness ranges from ∼80 to 160 m, is predominantly composed of mudstones and some interlayers of sandstones. This sedimentary body bears abundant plutonic and volcanic fragments from the lower Jurassic basement of the Honda Group (Saldaña Formation), as well as clasts of volcanic rocks formed in the magmatic arc of the Cordillera Central of Colombia during the Middle Miocene (Anderson et al., 2016).

**Figure 1.**
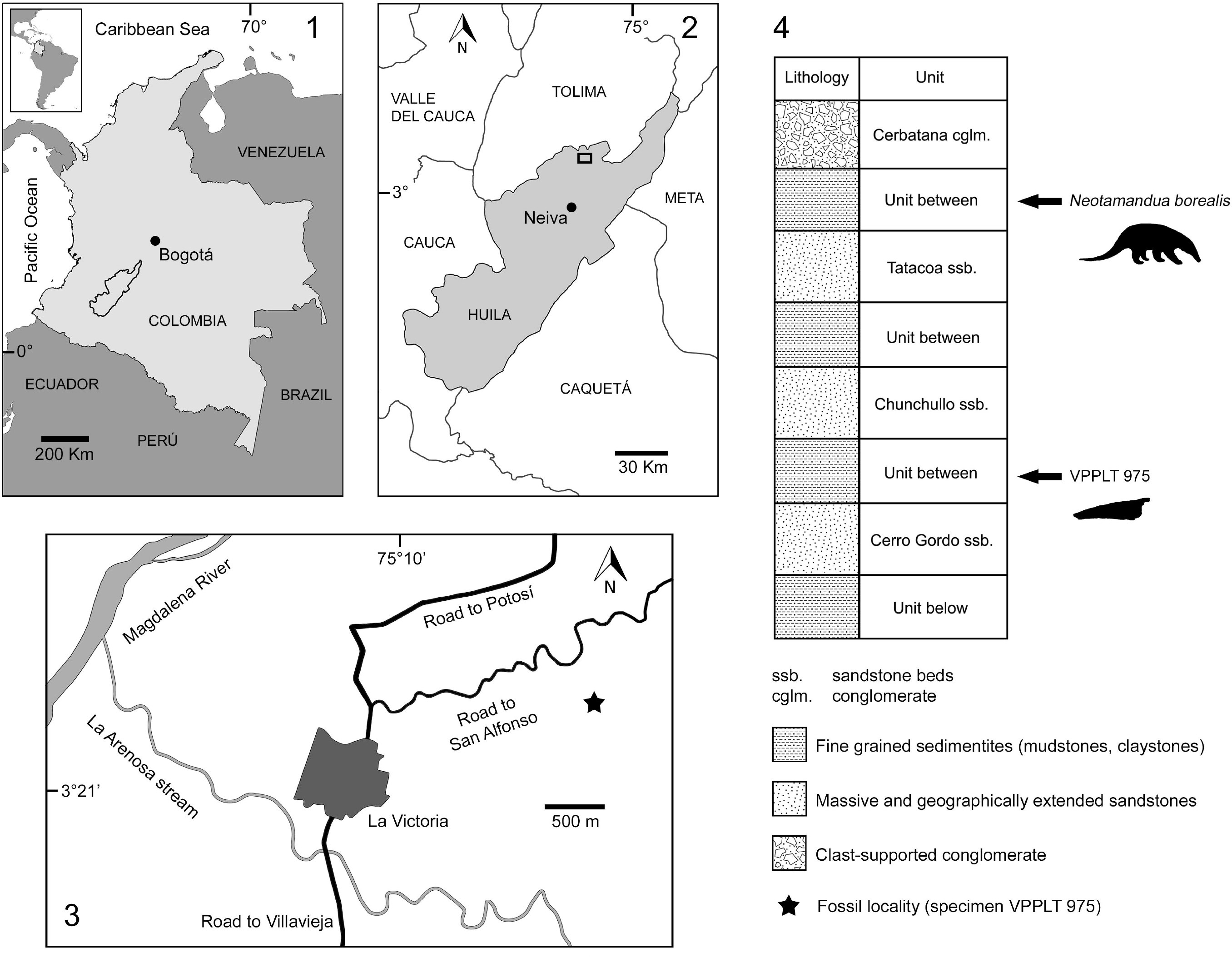
Geographic and stratigraphic provenance of the skull VPPLT 975 of the new taxon described here and the holotype of *Neotamandua borealis* (Hirschfeld, 1976). (**1**), location of the Department of Huila in Colombia; (**2**), location of the fossil area of interest, i.e., northern of La Venta area, in the Department of Huila (small rectangle); (**3**), location of the fossil site (black star), near the La Victoria town; (**4**), stratigraphic scheme of Guerrero (1997) for La Venta area, with approximate stratigraphic provenance of VPPLT 975 and the holotype of *N. borealis*.

The general paleoenvironment inferred for the La Victoria Formation is a meandering fluvial system (except for the Cerbatana conglomerate, associated to an anastomosed system) with significant soil development in flood plain zones (Guerrero, 1997). The ages calculated by Guerrero (1997) and Flynn et al. (1997) using magnetic polarity stratigraphy and geochronology, indicate sedimentary deposition took place in an interval of 13.8□12.5 Mya. These results have recently been reinforced by the U-Pb geochronology of detrital zircons recovered in this formation (Anderson et al., 2016), showing an age range of 14.4 ± 1.9 □ 13.2 ± 1.3 Mya. This interval coincides approximately with the early Serravalian, sub-stage of the Middle Miocene.

With some nomenclatural modifications, cranial measurements are based on those of Hossotani et al. (2017) (Fig. 2; see Anatomical Abbreviations). All these measurements are in millimetres (mm). The description of the new skull of La Venta includes a rough body mass estimation of the respective individual from a traditional allometric approach. All these data and analyses are compiled in the Supplementary material (Appendices S1 and S3). For the taxonomic analysis of the genus *Neotamandua*, the justifications of the generic allocations for the referred species (at least doubtfully) were revised in all the relevant scientific literature. These species are: *Neotamandua conspicua* Rovereto, 1914 (type species); *Neotamandua magna* Ameghino, 1919; *Neotamandua greslebini* Kraglievich, 1940; *Neotamandua borealis* Hirschfeld, 1976; *Neotamandua*? *australis* Scillato-Yané and Carlini, 1998. Additionally, some observations were made on the holotypes of *N. conspicua* (MACN 8097) and *N. borealis* (UCMP 39847) to reexamine the described characteristics for these species in the original publications (Rovereto [1914] and Hirschfeld [1976], respectively). The conceptual model of Plotcnick and Warner (2006) for the identification of taxonomic wastebaskets was applied to *Neotamandua*. From the foregoing and the designation of the specimen FMNH P14419 as epitype of *N. conspicua*, a diagnosis for *Neotamandua* was proposed. See a list of all the studied fossil specimens in the Appendix S1 of the Supplementary material.

**Figure 2.**
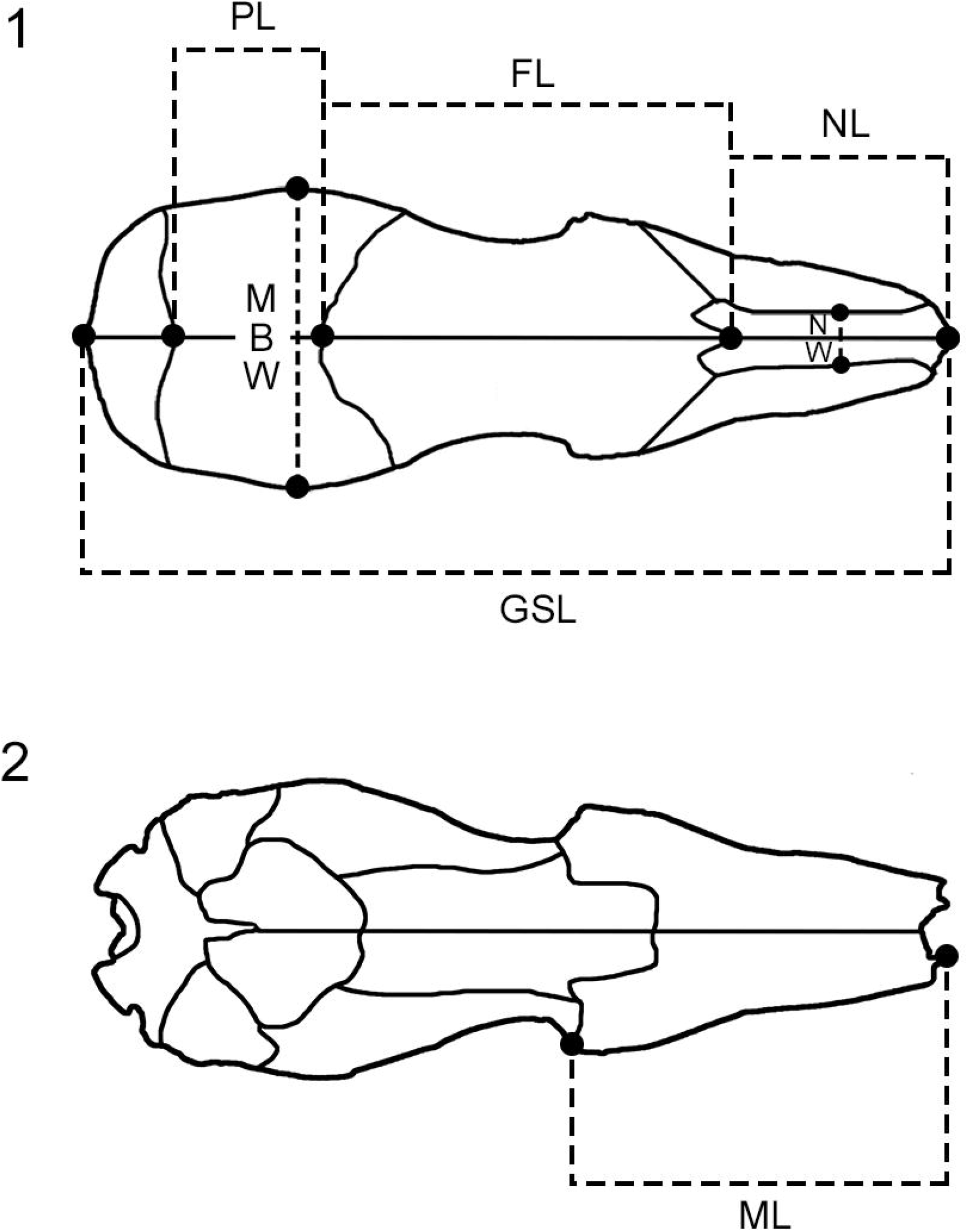
Cranial measurements used in this work. All of them are based on Hossotani et al. (2017). Nomenclatural modifications from these measurements are shown in the section of Anatomical Abbreviations. (**1**), skull of *Tamandua* in dorsal view; (**2**), the same skull in ventral view. *Abbreviations*. FL, frontal length; GSL, greatest skull length; MBW, maximum braincase width; ML, maxilla length; NL, nasal length; NW, nasal width; PL, parietal length.

**Figure 3.**
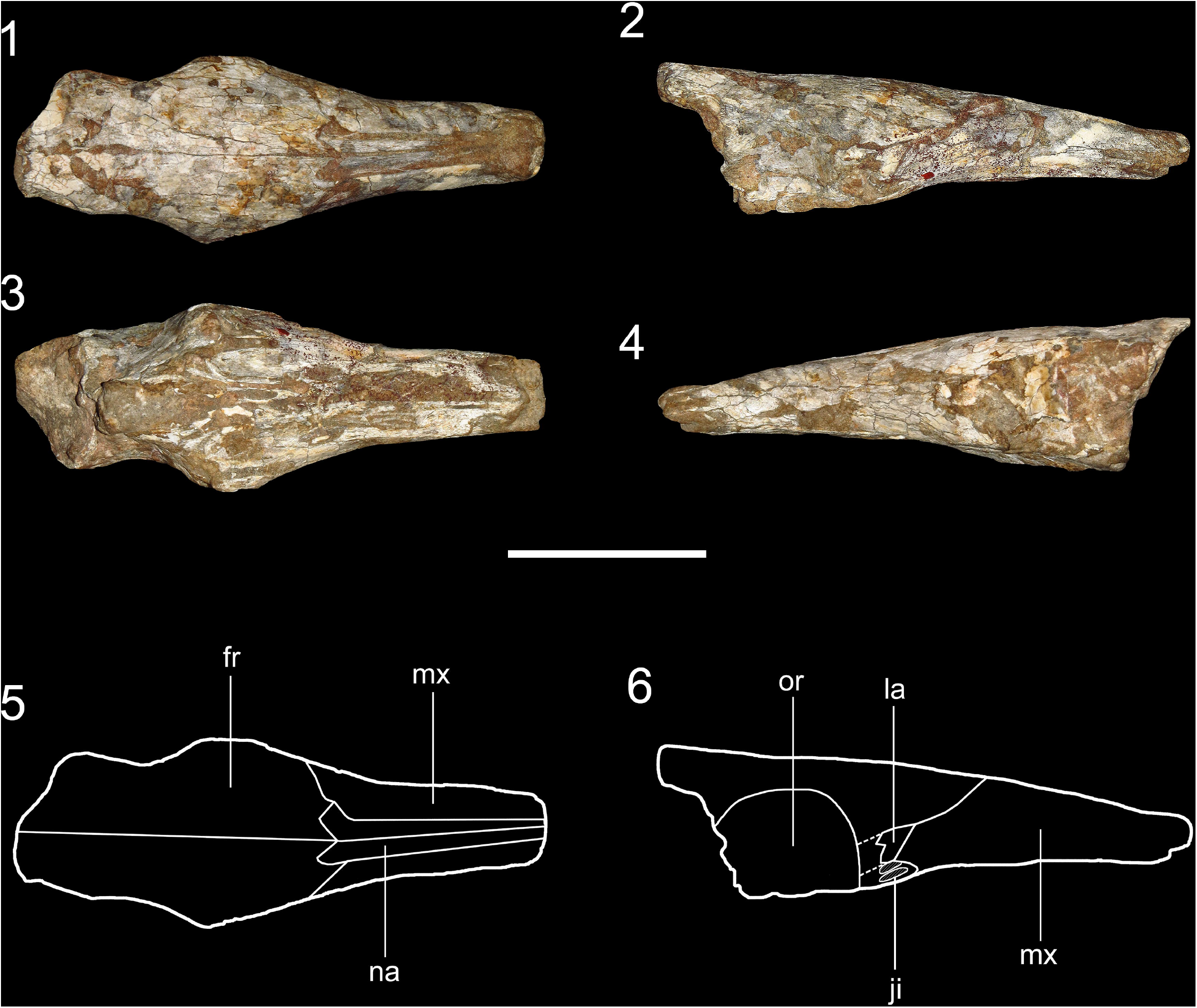
Holotypic skull (VPPLT 975) of Gen. et sp. nov. (**1**), dorsal view; (**2**), right lateral view; (**3**), ventral view; (**4**), left lateral view; (**5**), anatomical drawing in dorsal view; (**6**), anatomical drawing in right lateral view. *Abbreviations*. fr, frontal; ji, jugal insertion; la, lacrimal; mx, maxilla; na, nasal; or, orbit. Scale bar equal to 30 mm.

On the other hand, considering that Hirschfeld (1976), in her description of *N. borealis*, did not include morphological comparisons based on postcranial bones of this species and homologous elements of species referred to as *Neotamandua* from southern South America, we performed this task and a preliminary character distribution analysis from postcrania of these taxa to explore the hypothesis that they are closely related. Forcibly, *N. magna* and *N.*? *australis* are excluded from the comparisons since they do not have osteological elements correlated with those of *N. borealis* (or any other species referred to as *Neotamandua*). Additionally, as a result of the loss of its holotype (McDonald et al., 2008), comparisons with *N. greslebini* are based exclusively on the non-illustrated description by Kraglievich (1940). Other comparisons include postcranium collected by Juan Méndez in 1911 in the upper Miocene of the Andalhuala locality, Province of Catamarca, Argentina. This material was assigned to *Neotamandua* (*Neotamandua* sp.) without a reference publication. McDonald et al. (2008) manifested doubt about this taxonomic assignment (*Neotamandua*?), but these authors simultaneously speculated that it might be the lost holotype of *N. greslebini*.

Following McKenna and Bell (1997), the genus *Nunezia* is considered a junior synonym of *Myrmecophaga*. Myological inferences are based on Hirschfeld (1976) and Gambaryan et al. (2009).

### Repositories and institutional abbreviations

CAC: Cátedra de Anatomía Comparada, Facultad de Ciencias Naturales y Museo, Universidad Nacional de La Plata; FMNH: Field Museum, Chicago, IL., USA; ICN: Instituto de Ciencias Naturales, Facultad de Ciencias, Universidad Nacional, Bogotá, Colombia; MACN: Museo Argentino de Ciencias Naturales ‘Bernardino Rivadavia’, Buenos Aires, Argentina; MLP: Museo de La Plata, Facultad de Ciencias Naturales y Museo, Universidad Nacional de La Plata, La Plata, Argentina; MPT: Museo Provincial de Tucumán, Tucumán, Argentina; UCMP: University of California Museum of Paleontology, Berkeley, CA., USA; VPPLT: Museo de Historia Natural La Tatacoa, La Victoria, Huila, Colombia; YPM: Peabody Museum, Yale University, New Haven, CT, USA.

### Anatomical abbreviations

Abbreviations of equivalent measurements by Hossotani et al., (2017) in parenthesis. FL, frontal length; GSL (SL), greatest skull length; MBW (NC), maximum braincase width; ML, maxilla length; NL, nasal length; NW (NB), nasal width; PL, parietal length.

## Systematic paleontology

Superorder Xenarthra Cope, 1889

Order Pilosa Flower, 1883

Suborder Vermilingua Illiger, 1811

Family Myrmecophagidae Gray, 1825

New genus

### Type species

Gen. et sp. nov. (see below).

### Diagnosis

As for the type species by monotypy.

### Etymology

[this space is left in blank intentionally]

### Remarks

We propose the creation of Gen. nov. partly based on direct or indirect comparison between the holotypic skull of the only known and type species, Gen. et sp. nov., and cranial/postcranial material assigned to the species *Neotamandua borealis* and *N. conspicua* (see below the sections “Revised taxonomy of the genus *Neotamandua*” and “Systematic account for *Neotamandua*”).

Gen. et sp. nov.

### Holotype

VPPLT 975, anterior portion of a skull, without jugals nor premaxillae.

### Diagnosis

Middle sized myrmecophagid, slightly smaller than *Tamandua* and even more than *Neotamandua*. It can be differentiated from other genera/species of anteaters by the following combination of cranial features: relatively width rostrum, similar to *Tamandua*; narrow and strongly tapered nasals toward their anterior end; anteroposterior length of the preorbital section of frontals equal to more than two thirds of the anteroposterior length of nasals; jugals inserted from the same level of the most anterior border of the lacrimal; anterior portion of the orbit more laterally extended in the superior wall in the inferior one, without forming a conspicuous dome as in *Neotamandua conspicua*.

### Description

The rostrum is proportionally shorter and more robust than those in *Myrmecophaga* and *N. conspicua* (see below), but less than in *Tamandua*. In dorsal view, it is very similar to the skull of *Tamandua*, with at least four characteristics remarkably different with respect this extant genus: (1) lower rostrum; (2) rostrum more regularly tapered; (3) narrower and more anteriorly tapered nasals; (4) pre-orbital section of the frontals more anteroposteriorly elongated. In dorsal view, the rostrum shows a slight bulge in its middle part, similar to that in *Tamandua* and *Myrmecophaga*. However, in VPPLT 975 this bulge is even subtler than in the living myrmecophagids. Apparently, the nasals are shorter than frontals and are poorly exposed in lateral view. The jugals are absent by preservation, but it is possible to recognize their insertion location, which is more anterior than in *Myrmecophaga*, but more posterior than in *Tamandua*. Associated to the insertion of the jugal, there is a reduced posterolateral process of the maxilla in comparison with that of *Myrmecophaga*, similar in *Tamandua*. The right side of the skull preserves better the lacrimal zone, but it is simultaneously more deformed around the fronto-maxillary suture than in the left side. The lacrimal is longer in its anteroposterior axis than in that dorsoventral. The same bone is proportionally smaller than in *Tamandua* and even more than in *N. conspicua*. It has a triangular outline (at least anteriorly), similar to *Myrmecophaga* and unlike *Tamandua* (irregularly rounded, ovated, or, infrequently, sub-triangular lacrimal). The maxilla is not part of the orbit. The superior wall of the orbit is more laterally expanded than the inferior wall, without forming a conspicuous dome as in *N. conspicua*. This is similar to the condition observed in *Myrmecophaga* and differs from that in *Tamandua*, in which the inferior wall is prominent given that it is more laterally expanded. It is not possible to recognize lacrimal nor orbital foramina. In ventral view, the dorsal border of the orbit is regularly concave. The palatines are less laterally extended than in *Tamandua* and apparently there are no palatine “wings” (noticeable lateral expansions of the palatines), unlike *N. conspicua*.

### Etymology

[this space is left in blank intentionally]

### Remarks

See cranial measurements taken for this new species and other myrmecophagids in the Table 1. The estimated body mass for this individual is ca. 3.9 Kg (Appendix S3 of the Supplementary material). As consequence of the preservation, some sutures in VPPLT 975 are distinguishable in dorsal and lateral views, but virtually no suture is clearly detectable in ventral view.

**Table 1.**
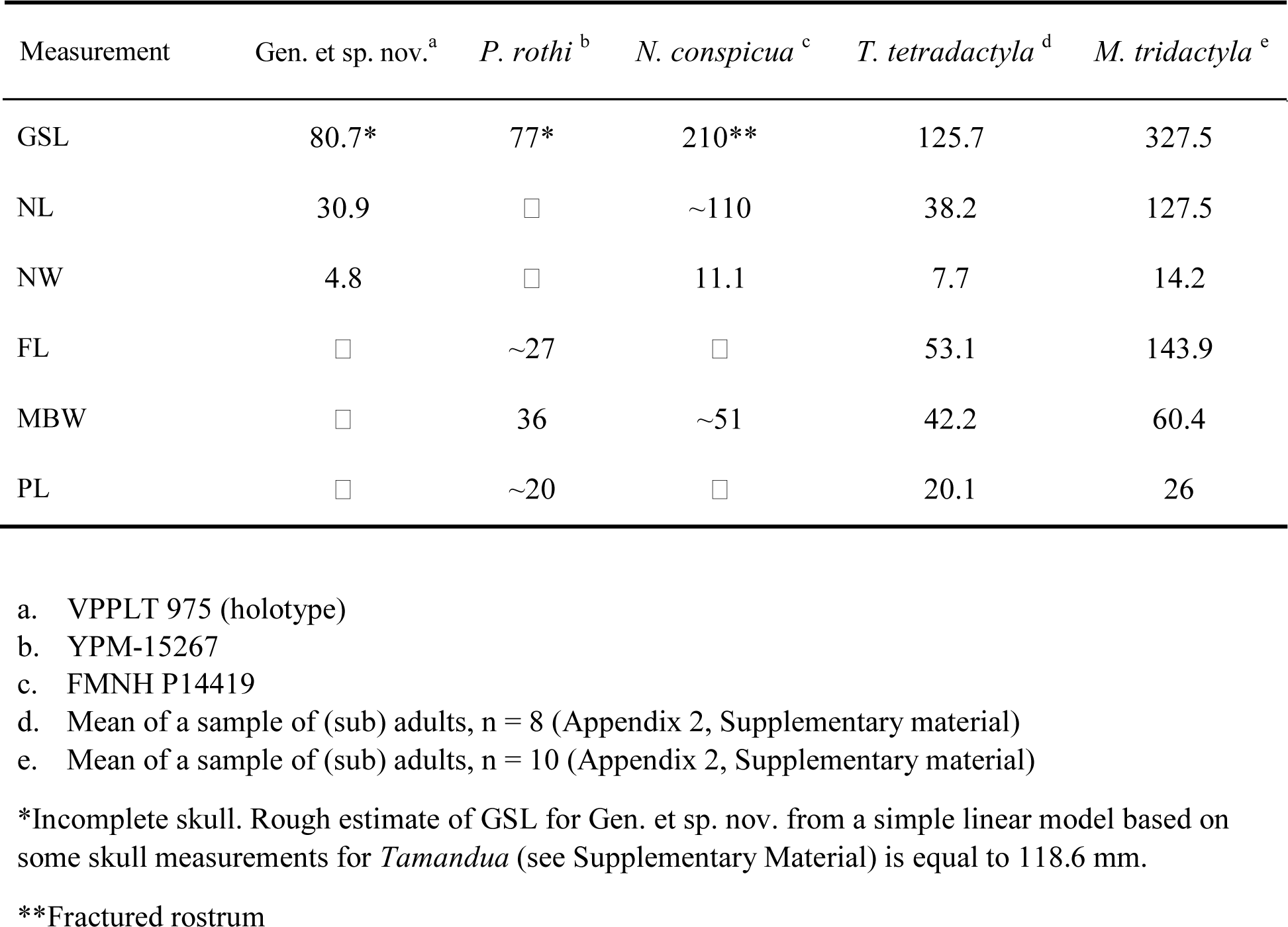
Cranial measurements (in mm) for the holotype of Gen. et sp. nov. and other myrmecophagid species.

## Revised taxonomy of the genus *Neotamandua*

### Taxonomic history and discussion on the taxonomic status of Neotamandua

The genus *Neotamandua* was proposed by Rovereto (1914) based on a posterior portion of a skull (MACN 8097), which was collected from upper Miocene-to-Pliocene strata of the Province of Catamarca, Argentina. The name *Neotamandua*, literally meaning ‘new tamandua’, was coined by Rovereto in allusion to the cranial similarity of the type species, *N. conspicua*, with the extant genus *Tamandua*, rather than with *Myrmecophaga*. This detail would be historically paradoxical, as will be shown below. It is important to note that Rovereto (1914) did not provide a diagnosis for *Neotamandua*, but he just briefly described the holotype of *N. conspicua*, emphasizing its elongated parietals. However, this feature, more comparable with that in *Myrmecophaga* than that in any other myrmecophagid, was correlated with the anteroposterior length of the parietals in *Tamandua*. A few years after Rovereto’s work, Carlos Ameghino (Ameghino, 1919) used a pelvis (MPT 58) recovered in contemporary strata from the Province of Tucuman, Argentina, to create a new species, *N. magna*. Despite the taxonomic assignment of this pelvis to *Neotamandua*, Ameghino discussed that, alternatively, this species could belong to other genus of larger body size, as Kraglievich (1940) also held. Formally, *N. magna* has not been reevaluated, but McDonald et al. (2008) suggested that, given that this species was transferred to *Nunezia* by Kraglievich (1934), and *Nunezia* is considered a junior synonym of *Myrmecophaga* (Hirschfeld, 1976; McKenna and Bell, 1997), then *N. magna* should be included in the latter genus, i.e., *Myrmecophaga magna* comb. nov. (unpublished). Indeed, the morphological differences cited by Ameghino (1919) and Kraglievich (1940) between the pelvis of *N. magna* and that of *M. tridactyla* (e.g., greater width and ventral flattening of the intermediate sacral vertebrae) do not seem sufficient to consider a generic distinction between these species.

Two decades later, Kraglievich (1940) proposed a new species based on postcranium collected in the upper Miocene of the Province of Catamarca. This was initially assigned to *N. conspicua*. According to Kraglievich, the then new species, *N. greslebini*, is easily identifiable by its large size, intermediate between those of *N. conspicua* and *N. magna*. Similar to Rovereto (1914), this author correlated with *Tamandua* his generic assignment of *N. greslebini* to *Neotamandua*, considering the similarity between the fossil specimens of this species and homologous elements of the living collared anteater (Kraglievich, p. 633). The holotype of *N. greslebini* is missing or mixed up with material labelled with generic names of extinct anteaters (i.e., *Neotamandua* and *Palaeomyrmidon*) in the Museo Argentino de Ciencias Naturales (MACN), in Buenos Aires, Argentina (McDonald et al., 2008).

Already in the second half of the 20^th^ century, a controversy about a possible synonymy between *Neotamandua* and *Myrmecophaga* arose. This means that there was a radical paradigmatic shift in myrmecophagid systematics from the early 20th-century perspectives which considered that *Neotamandua* is closely related to *Tamandua*, to the late 20^th^ century ones which considered that *Neotamandua* was even a serious candidate to be a junior synonym of *Myrmecophaga*. This historical change began with the non-cladistic systematic analysis of Hirschfeld (1976), in which *Neotamandua* was originally proposed as the direct ancestor (anagenetic form) of *Myrmecophaga*. In the same work, Hirschfeld created the first and, until now, only nominal extinct species of Vermilingua and Myrmecophagidae for northern South America, *N. borealis* (Middle Miocene of Colombia). Given the scarcity and fragmentation of the specimens referred to as *Neotamandua*, Hirschfeld (1976) recognized the need to revise the taxonomic validity of *N. conspicua, N. magna* and *N. greslebini*. Indeed, she went beyond and stated that *Neotamandua* species could be representatives of more than one single genus. However, her assignment of *N. borealis* to *Neotamandua* was based primarily on the idea that the fossils she studied are “considerably more advanced than those known from the Santacruzian [late Early Miocene], closer to the Araucanian [Late Miocene-Pliocene] species and…to the line leading to *Myrmecophaga* than *Tamandua*” (Hirschfeld, p. 421). For this author, many postcranial traits of *N. borealis* are intermediate between *Tamandua* and *Myrmecophaga*. As a questionable methodological aspect, it is important to note that Hirschfeld (1976) did not make osteological comparisons with the southern species of *Neotamandua*, only with postcrania of *Protamandua, Tamandua* and *Myrmecophaga* (extant species of the two latter genera).

In implicit reply to Hirschfeld (1976), Patterson et al. (1992) highlighted the morphological similarities between the unpublished skull FMNH P14419, catalogued as *N. conspicua* in the Field Museum, and the modern skulls of *Myrmecophaga*. For these authors, FMNH P14419 only differs from skulls of the living giant anteater in its smaller size. Consequently, Patterson et al. (1992) suggested to synonymize *Neotamandua* and *Myrmecophaga*, with nomenclatural priority for the latter. Nevertheless, Gaudin and Branham (1998) provided (weak) support for the validity of *Neotamandua* through a comprehensive phylogenetic analysis of Vermilingua. Their results indicate that *Neotamandua* is an independent taxon based on two autapomorphies, being one of them ambiguous and the other one unambiguous. The latter is the horizontal inclination of the glenoid. In the only most parsimonious tree recovered by Gaudin and Branham (1998), *Neotamandua* is closely related to *Myrmecophaga*, not *Tamandua*, as opposed to Rovereto (1914) and Kraglievich (1940).

Finally, the last species referred, with doubt, to the genus was *N.*? *australis* (Scillato-Yané and Carlini, 1998). The holotype of this species consists only of a humerus (MLP 91-IX-6-5) collected in the lower Middle Miocene of the Province of Río Negro, Argentina. Scillato-Yané and Carlini (1998) highlighted some similarities of this material with the humerus of *Tamandua*. They also expressed considerably uncertainty in assigning it to *Neotamandua*, not only by its fragmentary nature, but from the idea of Hirschfeld about the non-natural (i.e., non-monophyletic) status of this genus. Without performing a phylogenetic analysis, these authors proposed a hypothesis that *N. borealis* is closely related to *Myrmecophaga*, while *N. conspicua* and *N.*? *australis* are closer to *Tamandua*. If this hypothesis is correct, *N. borealis* does not belong to *Neotamandua* as consequence of the application of the nomenclatural priority principle.

In summary, multiple historical factors, including the lack of a diagnosis, insufficient number of anatomically correlatable/highly diagnostic postcranial elements and, especially, absence of cranial-postcranial associations, aroused the relatively arbitrary use of *Neotamandua* as a wastebasket taxon, i.e., a residual genus deriving from weak and/or inadequate systematic analysis. According to the conceptual model of Plotnick and Warner (2006), *Neotamandua* has five (from a total of seven) properties of a genus potentially classifiable as wastebasket: (1) it is an old name (i.e., more than one century to the present); (2) it is [relatively] rich in species (five species, i.e., the most speciose extinct genus of Vermilingua); (3) it has a [relatively] high number of occurrences; (4) it has wide temporal and geographic distributions; (5) it [primarily] groups together specimens poorly preserved and/or difficult to identify. To these five properties we may add the lack of a diagnosis, which is related in some way to the property number two of the Plotnick-Warner model, i.e., genera diagnosed from generalized characters, probably plesiomophies or easily recognizable characters.

As it was shown, *Neotamandua* has been invoked as a directly ancestral form of *Tamandua*, or, more recently, of *Myrmecophaga*, from its morphological characteristics in common with these two extant genera. But precisely because of this character mosaic, the generic allocation of isolated postcranial remains of myrmecophagids potentially referable to *Neotamandua* should not be reduced or exclusively focused on their comparison with the crown-group, but should also consider the effect of possible homoplasies (e.g., those related to ecological convergences), plesiomorphies and limitations of the fossil record (Plotnick and Warner, 2006). In other words, the apparent affinity between isolated postcranial elements of any Neogene anteater and their homologous in *Myrmecophaga* is not enough to make a reliable generic allocation in *Neotamandua*; diagnostic information of the latter genus is needed, preferably autapomorphies, which enable it to be individually identified and not simply as a set of forms similar to *Myrmecophaga*.

### Comparisons between northern and southern species referred to as Neotamandua

See the Table 2 and 3 for comparisons of postcranial measurements between these species.

**Table 2.**
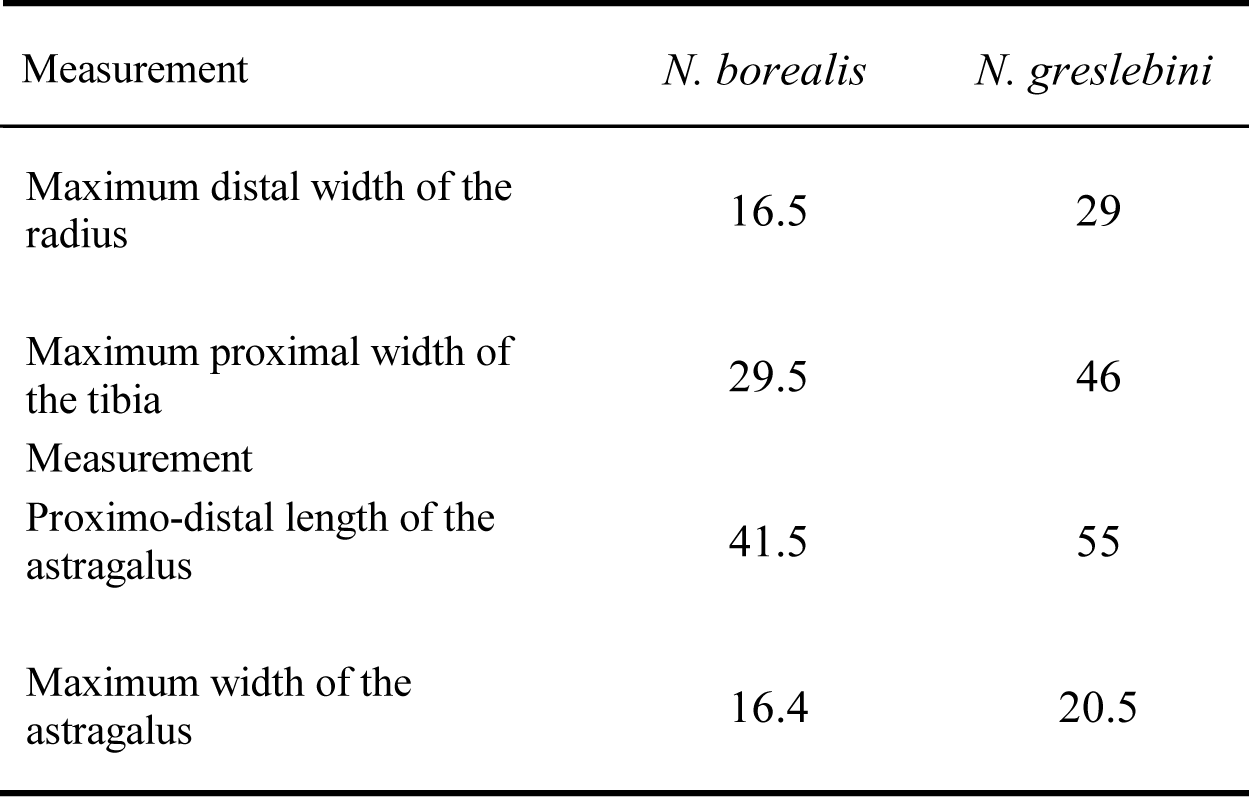
Comparison of some postcranial measurements (in mm) between *N. borealis* and *N. greslebini*. Data for the latter species after Kraglievich (1940).

**Table 3.** Comparison of some postcranial measurements (in mm) between *N. borealis* and *Neotamandua* sp.

|  |  |  |
| --- | --- | --- |
| Maximum distal width of the radius | 16.5 | 35 |
| Maximum width of the tibial articulation of the astragalus | 20 | 24 |
| Maximum width of the calcaneum | 20 | ~21 |

### N. borealis and N. greslebini

Both *N. borealis* and *N. greslebini* show two longitudinal, parallel radial ridges, of which the lateral ridge is higher and reaches a more distal level than the cranial one. This is similar to the condition observed in *Myrmecophaga* and differs from the distally convergent radial ridges of *Tamandua*. In *N. borealis*, the lateral ridge is even more distally extended than in *N. greslebini*, in such a way that the flanks of this structure contact the lateral border of the styloid process. According to Kraglievich (1940), in the Argentinean species this ridge ends at an intermediate level between the distal end of the cranial ridge and the styloid process.

The type material of *N. borealis* includes a proximal epiphysis and part of the diaphysis of a right tibia. According to Kraglievich (1940), the holotype of *N. greslebini* includes two fragments of a tibia, one of them proximal and the other one distal. Both Hirschfeld and Kraglievich claimed greater overall similarity between the tibial fragments of these species and the homologous parts of *Tamandua*, rather than *Myrmecophaga*. This way, the mid-section of the tibias both of *N. borealis* and *N. greslebini* is not as strongly triangular as in *Myrmecophaga*. Rather, this bone segment may vary from gently triangular to sub-rounded in these two species referred to as *Neotamandua*, not as rounded as in *Tamandua*.

Hirschfeld (1976) described the astragalus of *N. borealis* (Fig. 4) as intermediate in morphology and size between those in *Tamandua* and *Myrmecophaga*. In contrast, Kraglievich (1940) stated that the astragalus of *N. greslebini* closely resembles that of *Tamandua*. New observations enable us to determine that, in dorsal view, the astragalus of *N. borealis* is more similar to that in *Tamandua* than *Myrmecophaga* as consequence of a lateral side of the trochlea larger than the medial one (trochlear asymmetry). As *N. greslebini*, the regular concavity in which is inserted the *flexor digitorum fibularis* tendon extends posteroventrally like a well-defined wedge (“pointed shape” in Kraglievich’s words) and it contacts the calcaneal facets across the entire width of the latter. In ventral view, the arrangement of the calcaneal facets in *N. borealis* is a kind of “transition” between that in *Myrmecophaga* and *Tamandua*. In *N. borealis*, the ectal and sustentacular are largely separated by a wide and deep sulcus, but there is an incipient connection. This condition is approximately comparable to that described by Kraglievich (1940) for *N. greslebini* and differs from the fully separated calcaneal facets in *Protamandua* and *Tamandua*. In this sense, Kraglievich was not very explicit in pointing out the degree of development of the connection between these facets, but it is inferred that it is not exactly wide as in *Myrmecophaga* when he wrote that “…these calcaneal articulations are, *apparently*, posteriorly fused…” (italics are ours; Kraglievich, p. 635).

**Figure 4.**
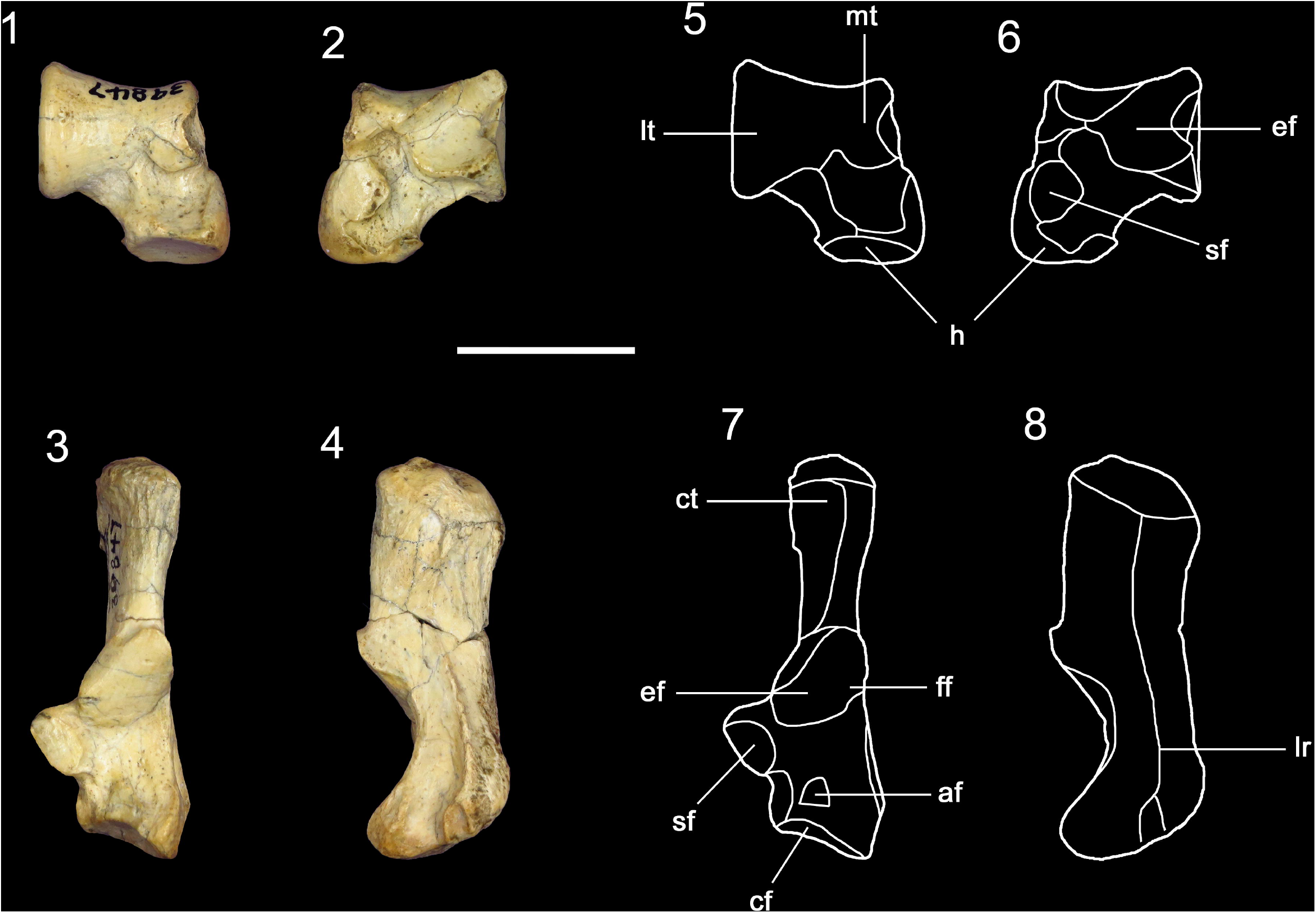
Two very informative postcranial bones of the holotype (UCMP 39847) of *Neotamandua borealis* (Hirschfeld 1976). (**1**), right astragalus, dorsal view; (**2**), right astragalus, ventral view; (**3**), left calcaneum, dorsal view; (**4**), left calcaneum, lateral view; (**5**), anatomical drawing of the astragalus in dorsal view; (**6**), anatomical drawing of the astragalus in ventral view; (**7**), anatomical drawing of the calcaneum in dorsal view; (**8**), anatomical drawing of the calcaneum in lateral view. *Abbreviations*. af, calcaneal accessory facet; cf, cuboid facet; ct, calcaneal tuber; ef, ectal facet; ff, fibular facet; h, astragalar head; lr, lateral ridge; lt, lateral trochlea; mt, medial trochlea; sf, sustentacular facet. Scale bar equal to 20 mm.

**Figure 5.**
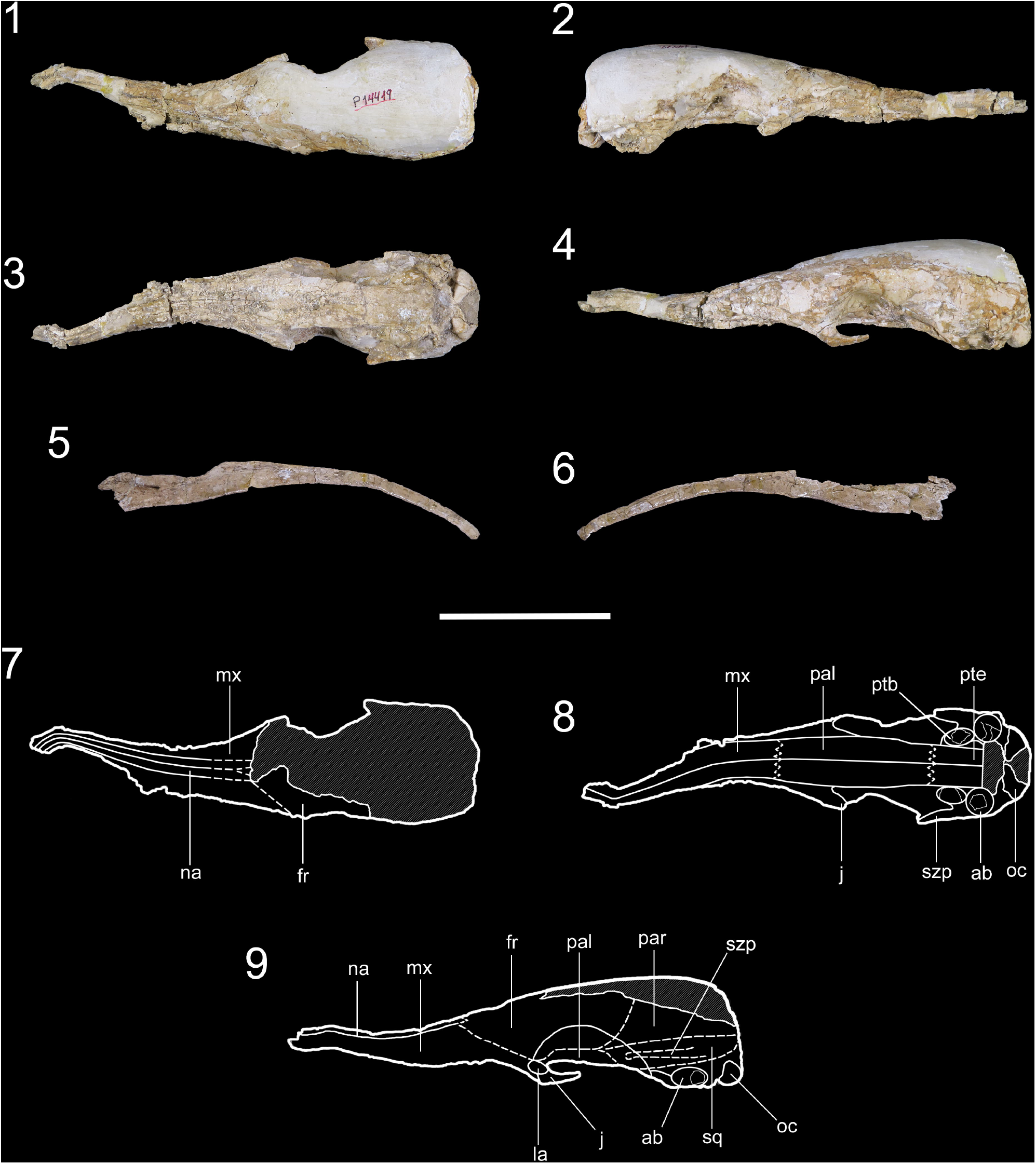
Epitype (FMNH P14419) of *Neotamandua conspicua*. (**1**), dorsal view; (**2**), right lateral view; (**3**), ventral view; (**4**), left lateral view; (**5**), right hemimandible; (**6**), left hemimandible; (**7**), anatomical drawing in dorsal view; (**8**), anatomical drawing in ventral view; (**9**), anatomical drawing in left lateral view. *Abbreviations*. ab, auditory bulla; fr, frontal; j, jugal; la, lacrimal; mx, maxilla; na, nasal; oc, occipital condyles; pal, palatine; par, parietal; ptb, pterygoid bulla; pte, pterygoid; sq, squamosal; szp, squamosal zygomatic process. Scale bar equal to 80 mm.

Similar to *N. greslebini, N. borealis* has a narrow fibular calcaneal facet, which is located laterally and in a slightly different plane with respect to that of the ectal facet (Fig. 4). In both of the former species, the *sustentaculum* is less medially projected than in *Myrmecophaga*. They also show an accessory facet in the anterior end of the calcaneum that articulates with the astragalar head, similarly to *Tamandua*. In all the aforementioned taxa, this facet is closer (even in contact) to the cuboid facet. In *N. borealis* and *N. greslebini*, the cuboid facet is transversely ovate and concave. A unique feature in common for them is the presence of a short tendinous groove (shorter than in *Myrmecophaga*) and strongly concave (Fig. 4). It is the continuation of the longitudinal and conspicuous ridge that runs the calcaneum in its lateral side. The latter separates tendons of the *fibularis longus* and *accesorius* muscles (Hirschfeld, 1976; Gambaryan et al., 2009). In *N. borealis*, this ridge is more conspicuous than in *Tamandua* and less than in *Myrmecophaga*.

### N. borealis and Neotamandua sp

See the Table 3 for comparison of postcranial measurements between these species. The distal epiphysis of the radius in *Neotamandua* sp. (MACN 2408) is more massive than that in *N. borealis*. In the latter species, the distal end of the radius is relatively stylized, similar to *Tamandua*. However, the morphologies of *N. borealis* and *Neotamandua* sp. are more comparable between them. In distal view, the styloid process of these species is more elongated and posteriorly oriented than in *Tamandua*. In the latter extant genus, the transverse axis (longer axis) of the facet for distal articulation is forming an angle close to 45° with respect to the plane of the anterior side of the radius, while this axis is nearly parallel with respect that plane in *N. borealis* and *Neotamandua* sp. This difference gives a non-rotated appearance to the distal radius of the compared *Neotamandua* species, unlike the same epiphysis in *Tamandua*. In anterior view, the distal articulation facet of *N. borealis* and *Neotamandua* sp. is visible in wedge shape pointing towards the medial border. Additionally, in the same view, this facet exhibits comparable exposures in both of the latter species, considerably more than in *Tamandua*. The posterior side of the distal epiphysis is from flat to slightly concave in *N. borealis* and *Neotamandua* sp., unlike the convex posterior side in *N. greslebini* (this observation could suggest that the material of *Neotamandua* sp. is not the holotype of *N. greslebini*, as speculated by McDonald et al., 2008) and *Tamandua*. The distal extension of the lateral ridge in *N. borealis* and *Neotamandua* sp. is similar.

The astragalus of *Neotamandua* sp. (MACN 2406) is only represented by the astragalar body. The medial trochlea is smaller than the lateral trochlea, but this asymmetry is lower than in *N. borealis*. In addition, these sections of the trochlea are proportionally less separated in the latter species than in *Neotamandua* sp. On the other hand, the calcaneum is fragmentary in *Neotamandua* sp. (MACN 2411). As in the case of the astragalus, the preserved portion is the bone body. The ectal facet is sub-triangular in shape in *Neotamandua* sp., while it is approximately sub-oval in *N. borealis*. The sustentacular facet is more medially extended in the latter species than in *Neotamandua* sp. In both species, the cuboid facet is partially visible in dorsal view, particularly in *Neotamandua* sp. In the same view, the lateral ridge is slightly exposed in *N. borealis*, not so much as in *Neotamandua* sp.

## Discussion

The former comparisons allow for recognizing a few morphological similarities and differences between homologous postcranial elements of *N. borealis, N. greslebini* and *Neotamandua* sp. It is considered that some similarities in these species are potentially diagnostic at the genus level, namely the sub-rounded to gently triangular shape of the tibial mid-section; ectal and sustentacular facets incipiently connected in the astragalus; and a short tendinous groove in the lateral side of the calcaneum (Table 4). These similarities seem to support the hypothesis that these northern and southern South American species referred to as *Neotamandua* are closely related and, consequently, that they are correctly included in the same genus. Alternatively, these common features could be symplesiomorphies of a hypothetical lineage of myrmecophagids more late diverging than *Protamandua* and apparently closer to *Myrmecophaga* than *Tamandua*. Provisionally, from the analysis presented, it is proposed to circumscribe the genus *Neotamandua* to the nominal species *N. conspicua* (type species), *N. greslebini* and *N. borealis*. Since *N. magna* and *N.*? *australis* are doubtfully assigned to *Neotamandua* or its allocation in this genus has been seriously questioned (McDonald et al., 2008; this work), they are considered *species inquirendae*, following the International Code of Zoological Nomenclature (Ride et al., 1999). To denote the questionable generic allocation of *N. magna* is suggested the use of inverted commas, i.e., ‘*N.*’ *magna*. The material referred to as *Neotamandua* sp. seems correctly referred to this genus, but it should be further tested. It is possible that these specimens correspond to a new species.

**Table 4.** Distribution of some postcranial characters of species referred to as *Neotamandua* and other myrmecophagid taxa. The characters marked with asterisk contain potentially diagnostic character states (synapomorphies?) for *Neotamandua*.

|  | *N. borealis* | *N. greslebini* | *Neotamandua* sp. | *Protamandua* | *Tamandua* | *Myrmecophaga* |
| --- | --- | --- | --- | --- | --- | --- |
| Relative body size | Medium to large | Large | Very large | Small | Medium | Very large |
| Character |  |  |  |  |  |  |
| 1. Arrangement of the radial ridges | Parallel ridges | Parallel ridges | Parallel ridges | NA | Distally convergent ridges | Parallel ridges |
| 2. Rotated appearance of the distal radius | Absent | Absent | Absent | Absent | Present | Absent |
| 3. Tibial mid-section* | Sub-rounded to gently triangular | Sub-rounded to gently triangular | NA | NA | Rounded | Strongly triangular |
| 4. Arrangement of the ectal and sustentacular facets in the astragalus* | Largely separated, but with an incipient connection | Largely separated, but with an incipient connection | NA | Fully separated | Fully separated | Widely connected |
| 5. Tendinous groove in the lateral side of the calcaneum* | Short | Short | NA | Absent | Absent | Long |

The diagnosis for *Neotamandua* proposed below is largely based on the designation of the specimen FMNH P14419 as epitype for the type species (see below), *N. conspicua*, after considering the fragmentary nature of the holotype of this taxon (MACN 8097; Rovereto, 1914), and, consequently, its ambiguity or lack of some taxonomically relevant features, particularly in the rostrum. In addition, the potentially diagnostic postcranial features for *Neotamandua* that has been identified above are also incorporated in the new diagnosis until cranial-postcranial associations are found and studied.

## Systematic account for *Neotamandua*

Suborder Vermilingua Illiger, 1811

Family Myrmecophagidae Gray, 1825

Genus *Neotamandua* Rovereto, 1914

### Type species

*N. conspicua* Rovereto, 1914.

### Other species

‘*N.*’ *magna* Ameghino, 1919; *N. greslebini* Kraglievich, 1940; *N. borealis* Hirschfeld, 1976; *N.*? *australis* Scillato-Yané and Carlini, 1998.

### Diagnosis

Middle-to-large sized myrmecophagid, larger than *Tamandua* but smaller than *Myrmecophaga*. It can be differentiated from other vermilinguans by the following combination of characteristics: in dorsal view, rostrum strongly tapered towards its anterior end (more than in any other myrmecophagid), with a regular transition in width from the anterior portion of the frontals to the anterior end of the nasals; reduced lacrimal which is not part of the orbit; jugal inserted in posteroventral position with respect to the lacrimal and slightly projected in posterodorsal direction; frontal forming a dorsal dome at the orbit level; hard palate well extended towards the posterior end of the skull, close to the ventral border of the occipital condyles; squamosal (= posterior) zygomatic process dorsally inclined; presence of palatine “wings”; horizontal inclination of the glenoid (Gaudin and Branham, 1998); sub-oval to gently triangular shape of the tibial mid-section; ectal and sustentacular facets incipiently connected in the astragalus; short tendinous groove in the lateral side of the calcaneum.

### LSID

urn:lsid:zoobank.org:act:4EC0ABE1-C013-4113-9956-5DBD6E79FCEA

*Neotamandua conspicua* Rovereto, 1914

### Holotype

MACN 8097, posterior portion of a skull.

### Epitype

FMNH P14419, nearly complete skull but with fractured rostrum and partially eroded frontals and parietals.

### Diagnosis

See the diagnosis for *Neotamandua* above. The postcranial diagnostic features included there do not belong to material known for this species.

### Occurrence

MACN 8097 is from an indeterminate locality in the Santa María Valley, Province of Catamarca, Argentina (Rovereto, 1914). Probably from the Andalhuala Formation. Upper Miocene (McDonald et al., 2008; Bonini, 2014; Esteban et al., 2014). FMNH P 14419 is from the Corral Quemado area, Province of Catamarca, Argentina. Corral Quemado Formation. Lower Pliocene (Bonini, 2014; Esteban et al., 2014). This specimen was collected by Robert Thorne and Felipe Méndez during the Second Captain Marshall Field Paleontological Expedition, which was led by Elmer S. Riggs and developed in Argentina and Bolivia in 1926□1927 (Simpson, pers. comm.; Riggs, 1928). In the Field Museum, where it is deposited, has been catalogued as *N. conspicua*. There are no known referencial publications that support the taxonomic assignation to this species, except in Gaudin and Branham (1998) and, now, in this work from direct comparison with its holotype.

### Description

The skull FMNH P14419 is anteroposteriorly elongated, with a general architecture which is more similar to that in *Myrmecophaga* than *Tamandua*. In dorsal view, both the rostrum, in general, as well as the nasals, in particular, are anteriorly tapered. The pre-orbital section of the frontals is proportionally less elongated than in *Myrmecophaga*. The lacrimal has a sub-triangular outline and its anteroposterior and dorsoventral lengths are similar, unlike *Myrmecophaga*, whose lacrimal is triangular and more anteroposteriorly elongated. The insertion of the jugals is more ventral and posterior than in *Myrmecophaga* and even more than *Tamandua*. Each jugal is slightly tapered by mediolateral compression in its posterior end and it is posterodorsally projected, instead of posteroventrally as in *Myrmecophaga*. The posterolateral process of the maxilla contacts the entire anterior and ventral borders of the lacrimal. The orbital ridge is less prominent than in *Myrmecophaga*. The superior orbital wall is laterally expanded, forming a roof more developed than in *Myrmecophaga*. At the orbit level, the palatines are also laterally expanded, forming palatine “wings”. These structures make the anterior hard palate look wider than the posterior palate. The posterior end of the hard palate is less ventrally projected, as opposed to *Tamandua* and *Myrmecophaga*. In lateral view, the squamosal zygomatic processes are dorsally inclined, contrary to the ventral inclination of the same bone projection in *Tamandua* and *Myrmecophaga*. This feature would be a convergence with *Cyclopes*. The braincase is proportionally larger than in *Myrmecophaga*, but smaller than in *Tamandua*. The tympanic bulla is less developed than in *Tamandua*. The external auditory meatus has subcircular to circular shape, similar to *Myrmecophaga* (ovated in *Tamandua*). In *N. conspicua* the same opening is located in a posterodorsal position, as *Myrmecophaga* and in contrast with *Tamandua*, in which it has an anterodorsal position. Despite the palatopterygoid suture is not well preserved, it appears to be more similar to the irregular suture in *Myrmecophaga*, with a posteriorly opened, asymmetrical “V” shape, than the regular suture in *Tamandua*, with an anteriorly opened, symmetrical “V” shape. There is no interpterygoid vacuity where a soft palate could be established, similar to *Myrmecophaga*. The occipital condyles are proportionally larger than in *Myrmecophaga*.

### LSID

urn:lsid:zoobank.org:act:C4DC62D5-6470-4A04-B152-D42ED3BA332C

### Remarks

The cranial measurements taken for FMNH P14419 are shown in the Table 1.

## Discussion

### Systematic implications

This works includes the first description of a new, valid extinct genus for Myrmecophagidae in the last century, i.e., Gen. nov. Likewise, it constitutes a novel taxonomic comprehensive reassessment for *Neotamandua* since the work by Hirschfeld (1976). The results suggest that there are still critical gaps in our knowledge on the composition and diversity of the Neogene assemblages of these xenartrans, particularly in the tropical region of South America. With the inclusion of Gen. et sp. nov. (Fig. 6), Myrmecophagidae now comprises at least five genera (three of them fully extinct) and 11 nominal species (eight extinct species), namely [the dagger means extinct species]: *Protamandua rothi*^†^; *Neotamandua*? *australis*^†^; *Neotamandua borealis*^†^; Gen. et sp. nov. ^†^; ‘*Neotamandua*’ *magna*^†^; *Neotamandua greslebini*^†^; *Neotamandua conspicua*^†^; *Myrmecophaga caroloameghinoi*^†^; *Myrmecophaga tridactyla*; *Tamandua tetradactyla*; and *Tamandua Mexicana* (McDonald et al., 2008; this work). Of these taxa, only two genera and two species have fossil occurrence in northern South America: *N. borealis* (Middle Miocene of Colombia; Hirschfeld, 1976) and Gen. et sp. nov. (Middle Miocene of Colombia; this work) (Fig. 7). The latter taxon is a small-to-middle sized myrmecophagid, comparable but slightly smaller than *Tamandua*. The general morphology of the skull of this new anteater resembles more to that of *Tamandua* than any other known taxon. It shows remarkable features such as: (1) strongly tapered nasals toward its anterior rostrum; (2) relatively low rostrum and anterior section of frontals; (3) large pre-orbital section of frontals; and (4) strongly triangulated (anterior) lacrimal. The tapering of nasals is a characteristic in common with *N. conspicua*, but in the latter species the entire rostrum is tapered, not only the nasals, as Gen. et sp. nov. The relatively low rostrum and anterior section of frontals seems to indicate a plesiomorphy, given that this feature is apparently present in *P. rothi*. A large pre-orbital section of frontals is shared, in (nearly) extreme condition, by *N. conspicua* and, especially, *Myrmecophaga*, but it should be noted that in Gen. et sp. nov. there is no such as elongated skull. And, finally, the strongly triangulated (anterior) lacrimal in the latter species is superficially similar to that in *Myrmecophaga*. Estimates of cranial measurements and features (rostrum length, exposure of the maxilla in the orbit and curvature of the basicranial-basifacial axis) used for coding the characters with numbers 4, 8, 9 and 42 of the character list by Gaudin and Branham (1998), enable us to tentatively infer the phylogenetic position of Gen. et sp. nov. as a taxon included within the clade *Tamandua* + *Neotamandua* + *Myrmecophaga* and located in a polytomy with *Tamandua*. Under this preliminary phylogenetic analysis, which is not presented in the results section because there is no enough information for coding the new taxon, *Protamandua* is well supported as the most basal myrmecophagid as consequence of sharing several character states with non-Myrmecophagidae Vermilingua (i.e., *Cyclopes* and *Palaeomyrmidon*; for more details, see Gaudin and Branham, 1998). For future studies, the subfamilial name “Myrmecophaginae” is tentatively suggested for all the Myrmecophagidae more late diverging than *Protamandua*, including possibly Gen. et sp. nov. In this sense, new and more complete material referable to the latter taxon is required to shed light on its phylogenetic position.

**Figure 6.**
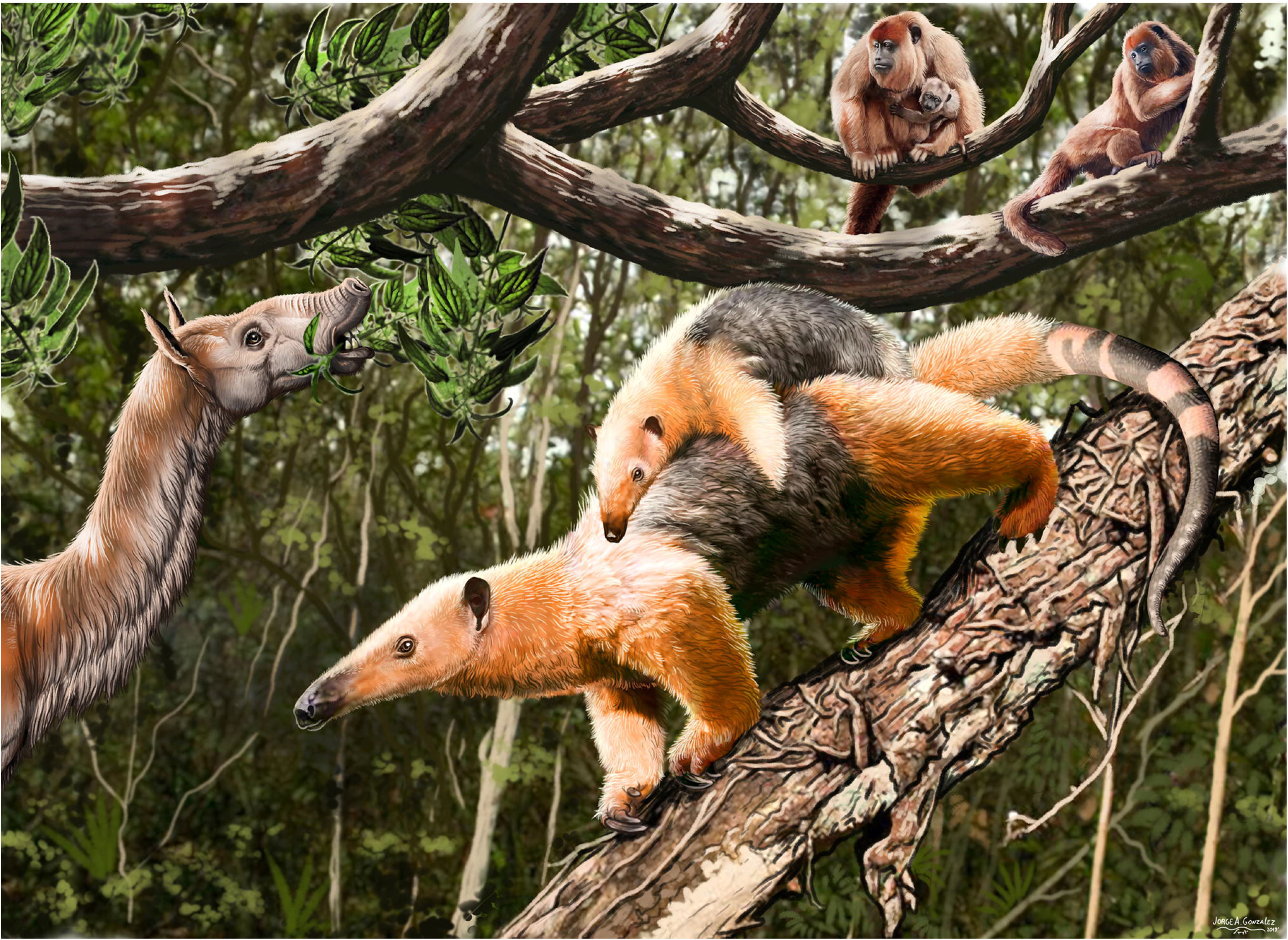
Reconstruction of the external appearance in life of Gen. et sp. nov. (close-up view). In the background, individuals of the macraucheniid *Theosodon* (left) and the alouattine *Stirtonia* (upper right corner) in the tropical forest of La Venta, late Middle Miocene of Colombia.

**Figure 7.**
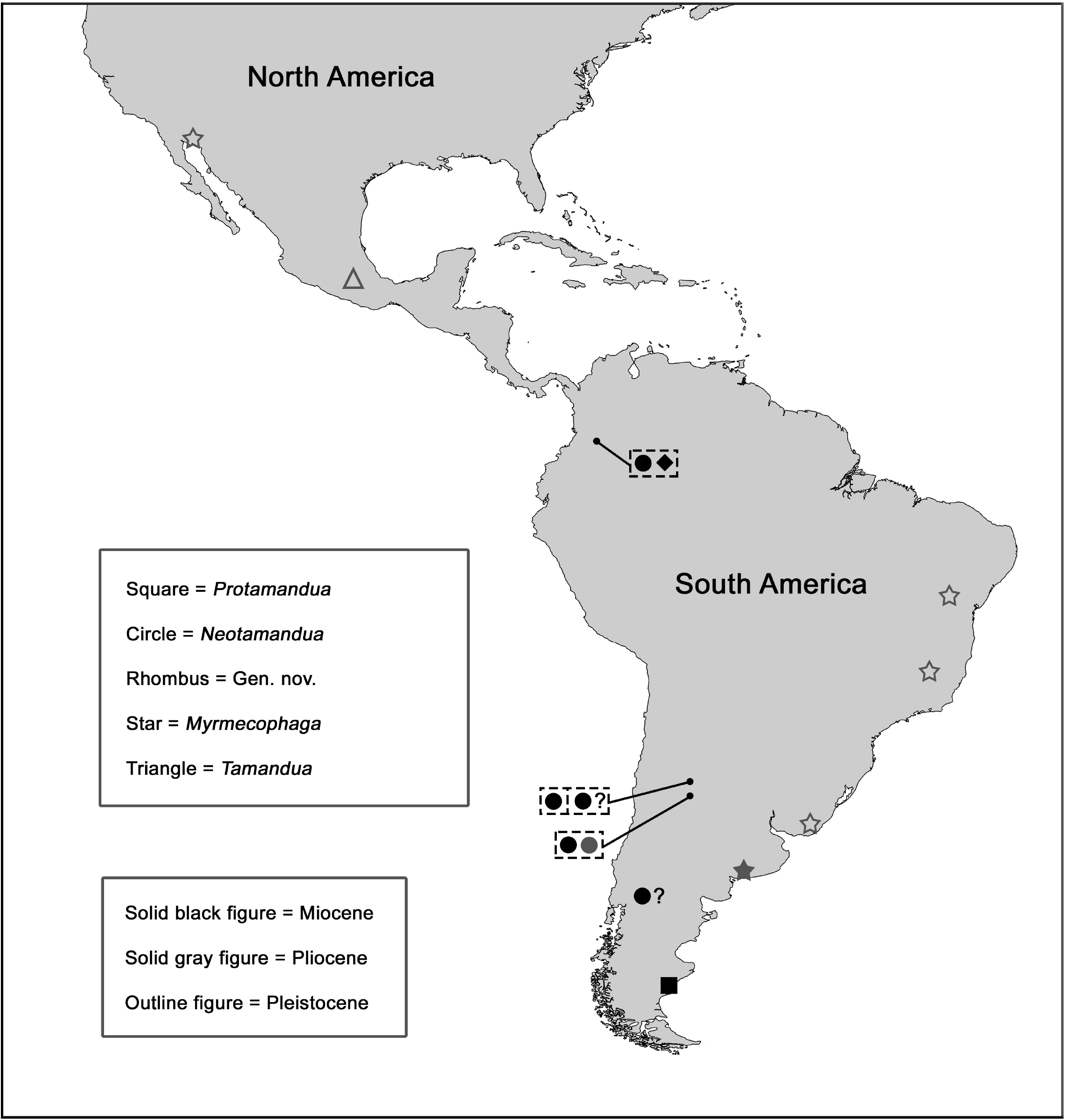
Geographic and chronological distribution of the myrmecophagid fossil records during the Late Cenozoic. Note the only two fossil records of these xenartrans outside South America in the Pleistocene of southern and northern Mexico (*Tamandua* sp. and *Myrmecophaga tridactyla*, respectively). Based on information compiled by McDonald et al. (2008). Original references in the same work and, largely, in the main text here.

On other hand, the taxonomic analysis of *Neotamandua* and its referred species indicates that these taxa were based on a poorly supported taxonomy. Other case of extinct vermilinguans with flawed systematics in low levels of the taxonomic hierarchy was noted by McDonald et al. (2008) with regard to genera and species proposed from isolated postcranial elements of putative myrmecophagids or even members of new, distinct families from the Early Miocene of Santa Cruz, southern Argentina. These authors, partially based on comparisons by Hirschfeld (1976), argued that the number of taxa claimed for that area and interval (seven genera and nine species; e.g., *Promyrmephagus, Adiastaltus*; Ameghino, 1894) has been artificially inflated, even though it is still possible to revalidate taxa other than the well validated species *P. rothi* (McDonald et al., 2008). All these research problems in systematics imply the need to regularly reevaluate the taxonomy of extinct anteaters through reexamination, when possible, of previously described material and the study of new specimens. While it is true that the fossil record of Vermilingua is poor and fragmentary in comparison, for instance, with that of other xenartrans such as the Tardigrada, the sampling effort should be increased in order to have greater recovery of fossil material for this group, especially in areas known for their preservation potential (e.g., southern and northwestern Argentina, southwestern Colombia).

Even though, no diagnosis for this genus was found through the reevaluation of the taxonomic status of *Neotamandua*. The newly proposed diagnosis includes multiple cranial and potential postcranial characteristics, which uphold that *Neotamandua*, whether it is a natural group or not, certainly contains species that do not belong to *Myrmecophaga*, despite their great resemblance with the latter. This outcome is congruent with the taxonomic opinion of Gaudin and Branham (1998) and is at odds with Patterson et al. (1992). Now, can we confidently say that *Neotamandua* is monophyletic from current evidence? As previously defined by other workers, *Neotamandua* may be composed of successive basal species or genera in relation to the hypothetical clade of *Myrmecophaga* (i.e., *My. tridactyla* + *My. caroloameghinoi*). If that is correct, *Neotamandua* would be paraphyletic by definition, since it excludes some of its descendants (Sereno et al., 1991). This possible pattern of basal paraphyly is consequence of a taxonomy that is not defined by clades, but grades (Huxley, 1958; Wood and Lonergan, 2008). The monophyly of *Neotamandua*, as was redefined here (i.e., *N. conspicua* + *N. greslebini* + *N. borealis*), is tentatively supported by three potential synapomorphies shared by two of its species whose postcranium is known (*N. greslebini* and *N. borealis*): (1) sub-oval to gently triangular mid-section of the tibia; (2) ectal and sustentacular facets incipiently connected in the astragalus; (3) short tendinous groove in the lateral side of the calcaneum. However, the synapomorphic conditions of these features for *Neotamandua* need to be further tested from systematic analysis of new, more complete and/or associated material of Gen. et sp. nov. and species referred to as *Neotamandua*. That would enable us to assess more adequately the global morphological variability and character distribution in Miocene myrmecophagids more late diverging than *Protamandua*. In turn, when there is a better knowledge of such distribution, it is more likely to disentangle the taxonomic identities and affinities of the *Neotamandua* species in order to corroborate the monophyly of this genus. For the moment, the hypothesis of Hirschfeld (1976) that *Neotamandua* is not monophyletic is, in principle, less probable if the *species inquirendae* ‘*N.*’ *magna* and *N.*? *australis* are excluded from the genus, as it was decided here, than if they are retained within it. The exclusion of the *species inquirendae* does not affect the hypothesis that *Neotamandua* is closer to *Myrmecophaga* than any other known nominal genus. Consequently, the type species of *Neotamandua, N. conspicua*, is reiterated as closer to *Myrmecophaga* than *Tamandua*, in line with the phylogeny of Gaudin and Branham (1998) and unlike the hypothesis of Scillato-Yané and Carlini (1998).

Finally, the material referred to as *Neotamandua* sp. that was used in this study to make comparisons with *N. borealis*, seems correctly allocated in that genus, but it might eventually be assigned to a new species with very large body size, larger than *N. greslebini*. This is partially conditioned to the clarification of the taxonomic status of ‘*N.*’ *magna*, which is a species comparable in body size to *Neotamandua* sp., so they could be (or not) the same taxon.

### The diversification of Myrmecophagidae

McDonald et al. (2008) pointed out that since the highly incomplete fossil record of Vermilingua, several fundamental questions on the evolution of this group, including morphological trends and the acquisition of ecological preferences in its distinct taxa, are largely unknown. Likewise, they highlighted some uncertainty related to the divergence times of possible sub-clades. However, several inferences and hypotheses about the evolutionary history of anteaters and, particularly, the myrmecophagids, can be outlined from the current evidence, including that presented in this work. Following Pascual and Ortiz-Jaureguizar (1990), McDonald et al. (2008) and Toledo et al. (2017), the next discussion is based on multiple paleobiological, ecological and biogeographic aspects as major constraints and/or consequences of the myrmecophagid evolution.

The diversification of Myrmecophagidae was a macroevolutionary event that occurred through the Neogene, at least as early as the Burdigalian (Early Miocene), according to the minimal age estimated for the most basal genus, i.e., *Protamandua*. The beginning of this diversification is approximately overlapped in time with the onset or development of similar events in other higher taxa in South America, such as the xenartrans Megatherioidea, Mylodontidae, Glyptodontidae and Dasypodini (Croft et al., 2007; McDonald and De Iuliis 2008; Bargo et al., 2012; Carlini et al., 2014; Boscaini et al., 2019), or the South American native ungulates Pachyrukhinae, Mesotheriinae and Toxodontidae related to *Pericotoxodon* and *Mixotoxodon* (Seoane et al., 2017; Armella et al., 2018a; Armella et al., 2018b). This pattern shows the importance of the Early Miocene, particularly the Burdigalian, as a critical interval for the diversification of multiple South American land mammal lineages. In light of the geographic provenance of *Protamandua*, the most probable ancestral area for Myrmecophagidae is southern South America (Fig. 7). The paleonvironmental conditions inferred for the Early Miocene of this area are considerably warmer and more humid (1000□1500 mm/year) than today, with presence of a subtropical dry forest (Iglesias et al., 2011; Quattrocchio et al., 2011; Kay et al., 2012; Brea et al., 2017; Raigenborm et al., 2018). In line with this reconstruction, Palazzesi et al. (2014) reported that southern Argentina harboured in the Early Miocene a plant richness comparable to that documented today for the Brazilian Atlantic Forest, in southeastern Brazil. Similar to *Tamandua, Protamandua* would have preferred forested habitats and would have had semiarboreal habits (Gaudin and Branham, 1998; McDonald et al., 2008; Kay et al., 2012). Whether the ancestral condition of substrate use in Myrmecophagidae is arboreal, as held by Gaudin and Branham (1998), the preference for open biomes (e.g., savannah) and terrestriality in *Myrmecophaga* (and possibly in *Neotamandua*) is a derived condition (McDonald et al., 2008; Toledo et al., 2017). The semiarboreal habits of *Tamandua* are explained from niche conservatism or, alternatively, from convergence with *Protamandua* if the ancestor of *Tamandua* was hypothetically terrestrial.

Since their particular, low basal metabolic rates and myrmecophagous diets (McNab, 1984, 1985), it is likely that the global warm recovery during the early Neogene (Early Miocene to early Middle Miocene; including the Middle Miocene Climatic Optimum or MMCO; Fig. 8), linked to a latitudinal temperature gradient reduction and an expansion of the tropical (warm) forest belt towards higher latitudes in the continents (including South America; see Anderson, 2009; Herold et al., 2011; Morley, 2011; Palazzesi et al., 2014), has influenced the evolutionary differentiation of the myrmecophagids, maybe predominantly *in situ* as in the climatically-induced evolution of other small Cenozoic mammals (Fortelius et al., 2014), such as *Protamandua*. This differentiation would have been triggered by an increase in the suitable area in terms of preferred biomes (warm forests in this case) and, especially, temporarily sustained availability of social insects for their feeding (McDonald et al., 2008; Kay et al., 2012; Toledo et al., 2017). Indeed, extant termites and ants (Termitidae and Formicidae, respectively) concentrate the vast majority of their biomass (and species richness) in the tropics and warm subtropical regions (Hölldobler and Wilson, 1990; Tobin, 1995; Davidson and Patrell-Kim, 1996; Eggleton et al., 1996; Davidson et al., 2003; Ellwood and Foster, 2004; Keller and Gordon, 2009). This ecogeographic pattern is consistent with the fossil record of the former higher taxa, which shows a strong tropical niche conservatism from their respective evolutionary origins in the Late Cretaceous/Early Paleogene (see below; Archibald et al., 2011; LaPolla et al., 2013; Guénard et al., 2015; Bourguignon et al., 2017).

**Figure 8.**
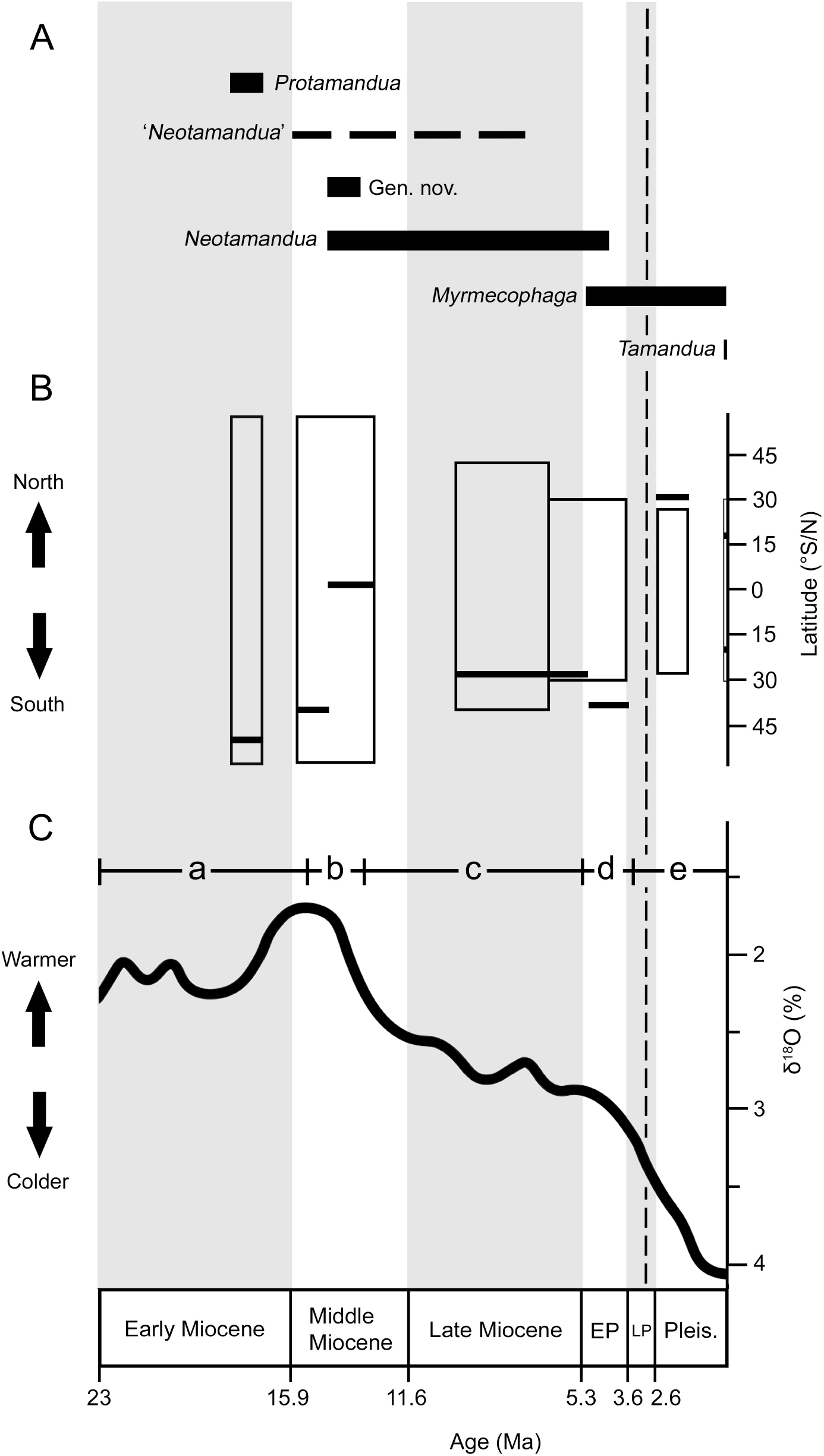
Chronological collation of data on: (**1**), biochrons of the myrmecophagid genera or questionable grouping (horizontal solid bars and dashed line, respectively); (**2**), distribution of the highest latitudinal fossil records (northern and/or southern) of myrmecophagids (horizontal solid bars) and approximate, chronologically discrete latitudinal ranges of tropical rainforest plus tropical and subtropical dry broadleaf forest (i.e., frost-free areas [mean annual temperatures higher than 15°C] with significant rainfall, at least seasonally; large vertical rectangles); (**3**), general trend curve of global temperature and climatic episodes during the Late Cenozoic: [a] early Neogene warm recovery, including the thermal peak in the late Early-early Middle Miocene known as Middle Miocene Climatic Optimum or MMCO; [b] Middle Miocene climatic transition; [c] late Middle-Late Miocene cooling; [d] Early Pliocene warming; [e] Late Pliocene-Pleistocene cooling and glaciations. The vertical dashed line indicates the time of complete formation of the Panama Land Bridge, which represented thereafter a fundamentally continuous physical connection between South- and North America. Paleocological data used for the plot in “B” is from the following references: Huntley and Webb III (1988); Toby Pennington et al. (2000); Williams et al. (2004); Williams (2009); Chan et al. (2011); Morley (2011); Kay et al. (2012); Pound (2012); Pound et al. (2012); Forrest et al. (2015); Lohmann et al. (2015); Dowsett et al. (2016); Sniderman et al. (2016); Henrot et al. (2017); Frigola et al. (2018). The temperature curve in “C” is based on Zachos et al. (2001, 2008) and it is reproduced with permission.

According to Blois and Hadly (2009), the responses of mammalian taxa to climate change throughout the Cenozoic are causally interconnected. These responses at the level of individual taxa may include changes in abundance, genetics, morphology and/or distributional range, and they may instigate multitaxa responses such as diversification events comparable to that placed on the root of the evolutionary tree of Myrmecophagidae. This case of a cladogenetic event possibly induced by climate contrasts in kind of biome with those that have been repeatedly documented for intervals of grassland expansion (e.g., Equidae, Bovidae, Cervidae, Ochotonidae, Hippopotaminae; MacFadden, 2000; Bouchenak-Khelladi et al., 2009; Boisserie and Merceron, 2011; Ge et al., 2013).

In the Middle Miocene, *N.*? *australis, N. borealis* and Gen. et sp. nov. exhibit a mosaic of morphological features in common with *Tamandua* and/or *Myrmecophaga*, as well as some exclusive characteristics, which suggest an early, important increase in morphological disparity in Myrmecophagidae and possibly the evolutionary divergence of those lineages comprising its crown-group. This coincides with the interpretation of Hirschfeld (1976), according to which the lineages that included the extant genera of Myrmecophagidae differentiated morphologically at least from the Friasian (Middle Miocene). Likewise, it is compatible with the results of the molecular phylogenies by Delsuc et al. (2001, 2012) and Gibb et al. (2016), which estimated that the evolutionary divergence of *Tamandua* and *Myrmecophaga* occurred in the late Middle Miocene, ca. 13 Mya. On the other hand, relative body sizes inferred for the Middle Miocene taxa show an apparent trend towards body size increase in comparison with the basal taxon *Protamandua*. During this interval, the myrmecophagids have a wide geographic distribution in South America (Fig. 7), from low to medium-high latitudes. This is in line with the evolution of larger body sizes since the increase in this attribute occurs, the foraging area also increases and, with it, the distributional range, according the general foraging strategy of the extant myrmecophagids (Naples, 1999; Toledo et al., 2017; Gaudin et al., 2018). The co-occurrence pattern of *N. borealis* and Gen. et sp. nov. in La Venta area in Colombia constitutes the earliest pattern of this kind for Myrmecophagidae until pending systematic revisions for putative taxa from the Early Miocene of Santa Cruz, Argentina, are carried out. These revisions would make it possible to determine whether there are two or more co-occurrent myrmecophagid taxa in the latter area. The fact that *N. borealis* and Gen. et sp. nov. are probably not sister taxa would imply a non-sympatric diversification followed by a dispersal of at least one of the involved taxa. The habitat preference of Gen. et sp. nov. in the paleoenvironmental mosaic of La Venta area (Kay and Madden, 1997; Spradley et al., 2019) is thought to be a tropical forest (semiarboreal?) by analogy with *Protamandua*, while it is proposed a more generalized habitat selection for *N. borealis* in line with the paleobiological inference of predominantly terrestrial locomotion for the latter taxon by Hirschfeld (1976). If this held true, it would open the possibility that *N. borealis* was the oldest myrmecophagid inhabiting zones with semi-open or even open vegetation (see below).

The morphological and probably taxonomic diversification of Myrmecophagidae continued in the Late Miocene. Inferred body sizes range from larger than *Tamandua* and nearly comparable to *Myrmecophaga*. Considering the wide geographic distribution during the Middle Miocene, there is probably a geographic bias in the fossil record of the myrmecophagids during the Late Miocene as the only known occurrences are *Myrmecophaga*-like forms from northwestern Argentina (Fig. 7). If *N. borealis* and *N. greslebini* are sister taxa, as it seems, that would mean that there was a biogeographic connection for Myrmecophagidae between northern and southern South America in the late Middle/early Late Miocene. This inference is congruent with the paleobiogeographic analyses of Cozzuol (2006) and Carrillo et al. (2015), according to which the affinities between several Late Miocene, northern and southern South American land mammal assemblages are strong or, at least, not so distant as those between Middle Miocene assemblages from the same regions. This pattern might be explained from the geographic shrinks of the Pebas Mega-Wetland System and the Paranean Sea in the Middle-Late Miocene transition (Aceñolaza and Sprechmann, 2002; Cozzuol, 2006; Salas-Gismondi et al., 2015). It is also possible that the expansion of open biomes in South America during the Late Miocene has facilitated this biotic connection, as it has been acknowledged in the case of other mammal taxa (e.g., Glyptodontinae, a xenartran group as Myrmecophagidae; Ortiz-Jaureguizar and Cladera, 2008; Oliva et al., 2010). Indeed, from a paleoenvironmental viewpoint, the (partial?) co-occurrence of ‘*N.*’ *magna, N. greslebini* and *N. conspicua* in northwestern Argentina is important inasmuch as this pattern is related, for the first time in the evolutionary history of Myrmecophagidae, to savannahs that were well developed with respect to other kinds of vegetation cover (Latorre et al., 1997; Brandoni et al., 2012; Cotton et al., 2014; Amidon et al., 2017; Zimicz et al., 2018). On the basis of the foregoing and by generalization of morphological and ecological features of the living vermilinguans, e.g., less dependence on trees related to greater taxonomic and/or ecological diversity of consumed insects (Hirschfeld, 1976; Montgomery, 1985a; Rodrigues et al., 2008; Toledo et al., 2017; Table 5), it is hypothesized that, as early as the late Middle Miocene, with the triggering of a global cooling (Fig. 8), *Neotamandua* was involved in a niche evolution process within Myrmecophagidae which implied a significative increase in dietary diversity as myrmecophagous and expansion of substrate use and biome selection. Probably the species of this genus preferred the frequent use of the ground by biomechanical constraints and made inroads into largely open environments as humid savannahs, without excluding the use of forested environments, as *Myrmecophaga* usually does (Fuster et al., 2018; Gaudin et al., 2018). The former model is further supported according to the evolutionary response pattern to major climatic-vegetational changes documented by Badgley et al. (2008) in a faunal sequence of land mammals from the Late Miocene of southern Asia, according to which the trophic niche evolution and, particularly the expansion of this attribute, in conjunction with habitat changes, are related to an increase in the probabilities of local and regional survivorship in the studied lineages.

**Table 5.** Taxonomic breadth in diet (genus level) of extant genera of Vermilingua and habitat preference of their MDCs (genera or species groups considered main dietary components). Key for the references: (1) Best and Harada (1985); (2) Fuster et al. (2018); (3) Gallo et al. (2017); (4) Gaudin et al. (2018); (5) Gómez et al. (2012); (6) Hayssen (2011); (7) Hayssen et al. (2012); (8) Jiménez et al. (2018); (9) Lubin and Montgomery (1981); (10) Medri et al. (2003); (11) Miranda et al. (2009); (12) Montgomery (1981); (13) Montgomery (1985a); (14) Montgomery (1985b); (15) Morales-Sandoval (2010); (16) Navarrete and Ortega (2011); (17) Redford (1985); (18) Rodrigues et al. (2008); (19) Sandoval-Gómez et al. (2012).

| Extant anteater genera | Termite genera | Ant genera | Some MDCs | Habitat preference of MDCs | References |
| --- | --- | --- | --- | --- | --- |
| *Cyclopes* | 0 | 8 | *Camponotus*<br>*Dolichoderus*<br>*Solenopsis* | Rainforest, seasonally dry tropical forest; trees | 1, 7, 11, 13, 19 |
| *Tamandua* | 7 | 17 | *Nasutitermes*<br>*Camponotus*<br>*Crematogaster*<br>*Solenopsis* | Seasonally dry tropical forest, rainforest; trees and ground | 3, 6, 9, 12, 14, 15, 16 |
| *Myrmecophaga* | 8 | 31 | *Cornitermes*<br>*Syntermes*<br>*Atta*<br>*Solenopsis*<br>*Camponotus*<br>*Acromyrmex*<br>*Pheidole* | savannah, grassland; ground | 2, 3, 4, 5, 8, 10, 14, 17, 18, 19 |

On the other hand, the fossil record of the crown-group genera, *Tamandua* and *Myrmecophaga*, is confined to the Pliocene-Pleistocene, but the evolutionary (morphological) divergence of *Myrmecophaga* would date back at least to the late Middle Miocene according the first appearance of *Neotamandua*, i.e., *N. borealis*. Under this assumption, the hypothesis of ‘*N.*’ *magna* as a species of *Myrmecophaga* is perfectly feasible. In any case, the biogeographic dynamics of the two extant myrmecophagid genera may have been constrained by their respective ecological tolerances and, these in turn, by the rapidly changing habitat and biome distribution in the Americas during at least the last five or six million years (de Vivo and Carmignotto, 2004; Salzmann et al., 2011; Sniderman et al., 2016; Amidon et al., 2017; Roberts et al., 2018; Grimmer et al., 2018). This applies especially to the case of *Tamandua* since this taxon is less generalist in relation to habitat selection than *Myrmecophaga* (McDonald, 2005). Considering the hypothesis of niche expansion for *Neotamandua*, the differentiation of *Myrmecophaga* would have accentuated this putative evolutionary trend through stronger preference for open environments, which is consistent with the general paleoenvironment of savannah in the Early Pliocene of the area where occurs the oldest species of the latter genus, i.e., *My. caroloameghinoi* (Zavala and Navarro, 1993; McDonald et al., 2008).

The myrmecophagid evolution has a late episode with the complete formation of the Panama Land Bridge (PLB) in the terminal Neogene (Coates and Stallard, 2013; O’dea et al., 2016; Jaramillo, 2018). *Myrmecophaga tridactyla* invaded and colonized Central- and southern North America (northern Mexico) at least as early as the Early Pleistocene (Shaw and McDonald, 1987; Fig. 7). This dispersal event is part of the Great American Biotic Interchange (GABI), specifically the episode referred as GABI 2 (Woodburne, 2010). Today, the northern boundary of this species is located in northern Central America, over 3000 Km to the south of the northernmost fossil record (Gaudin et al., 2018). This distributional difference was interpreted by Shaw and McDonald (1987) in the light of the occurrence of warmer and more humid conditions in the Early Pleistocene of southern North America (southern United States-northern Mexico) than today in the same area. These conditions would have enabled *Myrmecophaga* to colonize subtropical savannahs with permanent availability of insects included in its diet (Croxen III et al., 2007; McDonald, 2005). However, due extirpation, subsequent climatic-vegetational shifts (desertification) during the Late Pleistocene would have forced a range shrinkage of this taxon towards lower latitudes (McDonald, 2005; Ferrusquía-Villafranca et al., 2017). The distributional range pattern of tropical taxa expanded towards southern North America during some intervals of the Pleistocene has been well supported from the records of multiple taxa other than *Myrmecophaga*, including mammals and sauropsids (Shaw and McDonald, 1987; Moscato and Jasinski, 2016; Ferrusquía-Villafranca et al., 2017).

As *Myrmecophaga* did, *Tamandua* also colonized (or evolved in) northern continental territories outside South America. This is supported from the occurrence of *Tamandua* sp. in the terminal Pleistocene of Central Mexico (Arroyo-Cabrales et al., 2004; Ferrusquía-Villafranca et al., 2010; Fig. 7). In its northern zone, the current distributional area of *T. mexicana* includes latitudes comparable with that of the referred fossil record for this species (Navarrete and Ortega, 2011). Central Mexico is part of the transitional area between the current Neotropical and Neartic regions, called Mexican Transition Zone (MTZ; Halffter and Morrone, 2017). All these observations, in conjunction with the above interpretation of the Neogene biogeographic and environmental patterns, suggest that Myrmecophagidae kept throughout its evolutionary history a niche conservatism associated with tropical (warm) habitats (a case of phylogenetic niche conservatism or PNC; see Cooper et al., 2011; Fig. 8), possibly in parallel with the same pattern in species groups of its prey insects (Thompson, 1994). Even more, the fact that Myrmecophagidae currently accumulates its highest species richness in the warmest and wettest belt of the Americas (Hayssen, 2011; Navarrete and Ortega, 2011; Miranda et al., 2017; Gaudin et al., 2018) is further interpreted as evidence that this higher taxon represents support for the tropical niche conservatism hypothesis (TCH; Wiens and Donoghue, 2004; Wiens and Graham, 2005). However, in line with the discussion above, this major ecological constraint in Myrmecophagidae is not only related to environmental thermal tolerance (see McNab [1985] for an analysis on the thermophysiological constraints of the Xenarthra; McNab [1984] also discussed the same issue for myrmecophagous mammals), as emphasized by TCH, but it is also driven by food availability, at least by limiting or preventing historical colonization of low-productivity regions far from the tropics (Shaw and McDonald, 1987; McDonald, 2005; Šímová and Storch, 2017; Fig. 8).

## Conclusion

The systematic evidence presented here suggests that probably the diversification of Myrmecophagidae is taxonomically and biogeographically more complex than previously thought. This insight is based on the description of the new taxon Gen. et sp. nov. for the Middle Miocene of Colombia (co-occurrent species of *N. borealis*) and the determination of *Neotamandua*, as previously defined, as a wastebasket taxon which is probably formed by species belonging to more than one single genus. While Gen. et sp. nov. possibly has affinities with *Tamandua*, more information is needed to test its phylogenetic position within Myrmecophagidae. On the other hand, *N. borealis, N. greslebini* and *Neotamandua* sp. share postcranial features (potential synapomorphies) that imply some grade of kinship between them. Therefore, the two nominal species among the former ones are provisionally kept within *Neotamandua*. Alternatively, these features may also constitute symplesiomorphies of a hypothetical lineage which is apparently close to *Myrmecophaga*. The remaining nominal species referred to as *Neotamandua*, i.e., ‘*N.*’ *magna* and *N.*? *australis* were designated as *species inquirendae*. Overall, it is necessary to develop new systematic revisions, including new phylogenetic analyses similar to that by Gaudin and Branham (1998), using new material referable to Gen. et sp. nov. and the referred species to *Neotamandua*, so as to obtain enough evidence to solidly determine the phylogenetic position of the new species from La Venta and corroborate the putative monophyletic status of *Neotamandua*. In line with the foregoing considerations, the paleontological exploration of Neogene sedimentary units in northern South America and northern Argentina is crucial to improve our understanding of the diversification of Myrmecophagidae.

## Acknowledgements

A lot of people and institutions have contributed in distinct ways to the development of this research. We would like to specially thank A. Vanegas and his family; the group of defenders of the paleontological heritage *Vigías del Patrimonio Paleontológico La Tatacoa* (Colombia); C. Jaramillo (Smithsonian Tropical Research Institute, Panama); and all the staff of the Museo de Historia Natural La Tatacoa (Colombia) for assistance and providing access to the holotype of the new species described for La Venta. We thank A. Kramarz (MACN, Argentina), K. Angielczyk (FMNH, United States), W. Simpson (FMNH, United States) and P. Holroyd (UCMP, United States), E. Flórez (ICN, Colombia), A. Carlini (CAC, Argentina) and I. Olivares (MLP, Argentina) gave permission to study extinct and extant anteater specimens in collections under their care. We are also grateful to S. Vizcaíno and S. Bargo (MLP – CONICET, Argentina) as they provided photos of postcranial specimens labelled as *Neotamandua* sp. in the MACN; D. Brinkman and J. Henderson (YPM, United States) shared photos of a skull of *Protamandua rothi* (YPM-15267); M. Pound (Northumbria University, United Kindom) and C. Hossotani (Universidade Federal de Mato Grosso do Sul, Brazil) kindly helped by facilitating their postgraduate theses; and J. Zachos (University of California – Santa Cruz, United States) gave permission to reuse part of the data for his Cenozoic global temperature curve. Finally, J. Gonzalez was the paleoartist who exquisitely recreated the new anteater from La Venta. Financial support given by the following institutions: CONICET (Internal Doctoral Fellowship); Field Museum (Science Visiting Scholarship); University of California Museum of Paleontology (Welles Fund).

## Statement of data archiving

The nomenclatural acts contained in this work are registered in Zoobank: [identifier Gen. et sp. nov.]

*LSID.* urn:lsid:zoobank.org:act:4EC0ABE1-C013-4113-9956-5DBD6E79FCEA

*LSID.* urn:lsid:zoobank.org:act:C4DC62D5-6470-4A04-B152-D42ED3BA332C

## Accesibility of supplemental data

Data available from the Dryad Digital Repository: [intentional blank space]

