## Supplementary material for "The first fossil skull of an anteater (Vermilingua, Myrmecophagidae) from northern South America, a taxonomic reassessment of *Neotamandua* and a discussion of the myrmecophagid diversification"

^2^CONICET, Consejo Nacional de Investigaciones Científicas y Técnicas, Argentina.

^3^Independent researcher, Buenos Aires, Argentina <>

**Appendices**

**S1.** List of specimens or samples of the studied taxa

**S2.** Distribution of values for cranial measurements in (sub) adult samples of *Tamandua tetradactyla* (n = 8) and *Myrmecophaga tridactyla* (n = 10)

**S3.** Data and regression analyses to estimate greatest skull length (GSL) and body mass for the specimen VPPLT 975, referred to Gen. et sp. nov.

**S1.** List of specimens or samples of the studied taxa

| **Taxon** | **Specimen/sample catalogue number** | **Geographical provenance** |
| --- | --- | --- |
| Gen. et sp. nov. | VPPLT 975 | Huila, Colombia |
| *Protamandua rothi* | YPM-15267, FMNH P13134 | Santa Cruz, Argentina |
| *Neotamandua conspicua* | MACN 8097, FMNH P14419 | Catamarca, Argentina |
| *Neotamandua* *borealis* | UCMP 39847 | Huila, Colombia |
| *Neotamandua* sp. | MACN 2403, 2406, 2408, 2411 | Catamarca, Argentina |
| *Tamandua tetradactyla* | ICN 487, 760, 3720; MLP 615, 1233, 2216, 8-IX-98-1; CAC 49254 | Meta, Colombia; Misiones, Argentina; indeterminate provenance in Argentina |
| *Tamandua mexicana* | ICN 12979, 21215 | Boyacá and Santander, Colombia |
| *Myrmecophaga tridactyla* | ICN 715, 734, 1041, 21316, 21465; MLP 140, 15-III-51-1; CAC 62, 138, CAC indeterminate number | Meta and Casanare, Colombia; indeterminate provenance in Argentina |

**S2.** Distribution of values for cranial measurements in (sub) adult samples of *Tamandua tetradactyla* (n = 8) and *Myrmecophaga tridactyla* (n = 10).

| Species | Specimens | GSL | NL | NW | FL | MBW | PL |
| --- | --- | --- | --- | --- | --- | --- | --- |
| *Tamandua tetradactyla* | ICN 487 | 117.29 | 38.33 | 7.61 | 48.19 | 43.29 | 19.52 |
|  | ICN 760 | 126.56 | 40.69 | 7.94 | 53.1 | 40.38 | 22.43 |
|  | ICN 3720 | 130.21 | 42.58 | 6.76 | 55.43 | 42.71 | 21.47 |
|  | MLP 615 | 110.89 | 31.58 | 7.09 | 52.52 | 38.43 | 14.84 |
|  | MLP 1233 | 119.01 | 31.36 | 7.7 | 53.05 | 41.16 | 17.7 |
|  | MLP 2216 | 132.51 | 39.47 | 8.1 | 52.52 | 44.82 | 22.71 |
|  | MLP 8-IX-98-1 | 135.08 | 46.69 | 8.4 | 49.7 | 42.04 | 18.67 |
|  | CAC 49254 | 134.53 | 35.07 | 8.6 | 60.57 | 44.61 | 23.7 |
| *Myrmecophaga tridactyla* | ICN 715 | 358.97 | 137.56 | 14.55 | 176.45 | 56.17 | 21.43 |
|  | ICN 734 | 337.39 | 116.74 | 16.4 | 154.06 | 56.42 | 28.73 |
|  | ICN 1041 | 330.87 | 129.6 | 14.66 | 147.74 | 61.05 | 22.14 |
|  | ICN 21316 | 342.8 | 131.78 | 14.12 | 156.25 | 56.52 | 23.6 |
|  | ICN 21465 | 334.3 | 143.8 | 14.7 | 155.9 | 61.7 | 25.15 |
|  | MLP 140 | 314 | 112.69 | 13.5 | 135.98 | 59.40 | 30.39 |
|  | MLP 15-III-51-1 | 371 | 157 | 15.8 | 150 | 65.88 | 24.09 |
|  | CAC 62 | 308 | 119.2 | 13.4 | 128.35 | 63.24 | 29.9 |
|  | CAC 138 | 330 | 136.57 | 14.1 | 134 | 67.64 | 24.75 |
|  | CAC indeterminate number | 248 | 90.49 | 10.7 | 99.88 | 55.3 | 29.44 |

**S3.** Data and regression analyses to estimate greatest skull length (GSL) and body mass for the specimen VPPLT 975, referred to Gen. et sp. nov.

All cranial measurements are in millimetres (mm) and body mass in grams (g). A predictive equation derived from a least-squares regression for the values of maxilla length (ML) and greatest skull length (GSL) from an unpublished dataset of 10 (sub) adults of *Tamandua tetradactyla* (eight) and *T. mexicana* (two) was used to approximately estimate the GSL of the new fossil anteater. Given that no body mass data were available for these individuals whose cranial measurements were known, a consecutive regression was run to model body mass *versus* GSL from a distinct, published dataset (n =10) of *T. tetradactyla* from Suriname, northeastern South America (Husson 1978). In the former regression analysis, all measurements were log-transformed in advance for the purpose of diminishing any major difference in variance between the variables.

**Table 1.** Unpublished dataset on the distribution of maxilla length (ML) and greatest skull length (GSL) in 10 specimens of *Tamandua* from Colombia and Argentina.

| **Taxon** | **Specimen** | **ML** | **GSL** |
| --- | --- | --- | --- |
| *Tamandua tetradactyla* | ICN 487 | 48.49 | 117.29 |
|  | ICN 760 | 57.12 | 126.56 |
|  | ICN 3720 | 57.52 | 130.21 |
|  | MLP 615 | 46.96 | 110.89 |
|  | MLP 1233 | 49.05 | 119.01 |
|  | MLP 2216 | 56.32 | 132.51 |
|  | MLP 8-IX-98-1 | 58.97 | 135.08 |
|  | CAC 49254 | 55.84 | 134.53 |
| *Tamandua mexicana* | ICN 12979 | 58.76 | 124.65 |
|  | ICN 21215 | 57.46 | 122.78 |

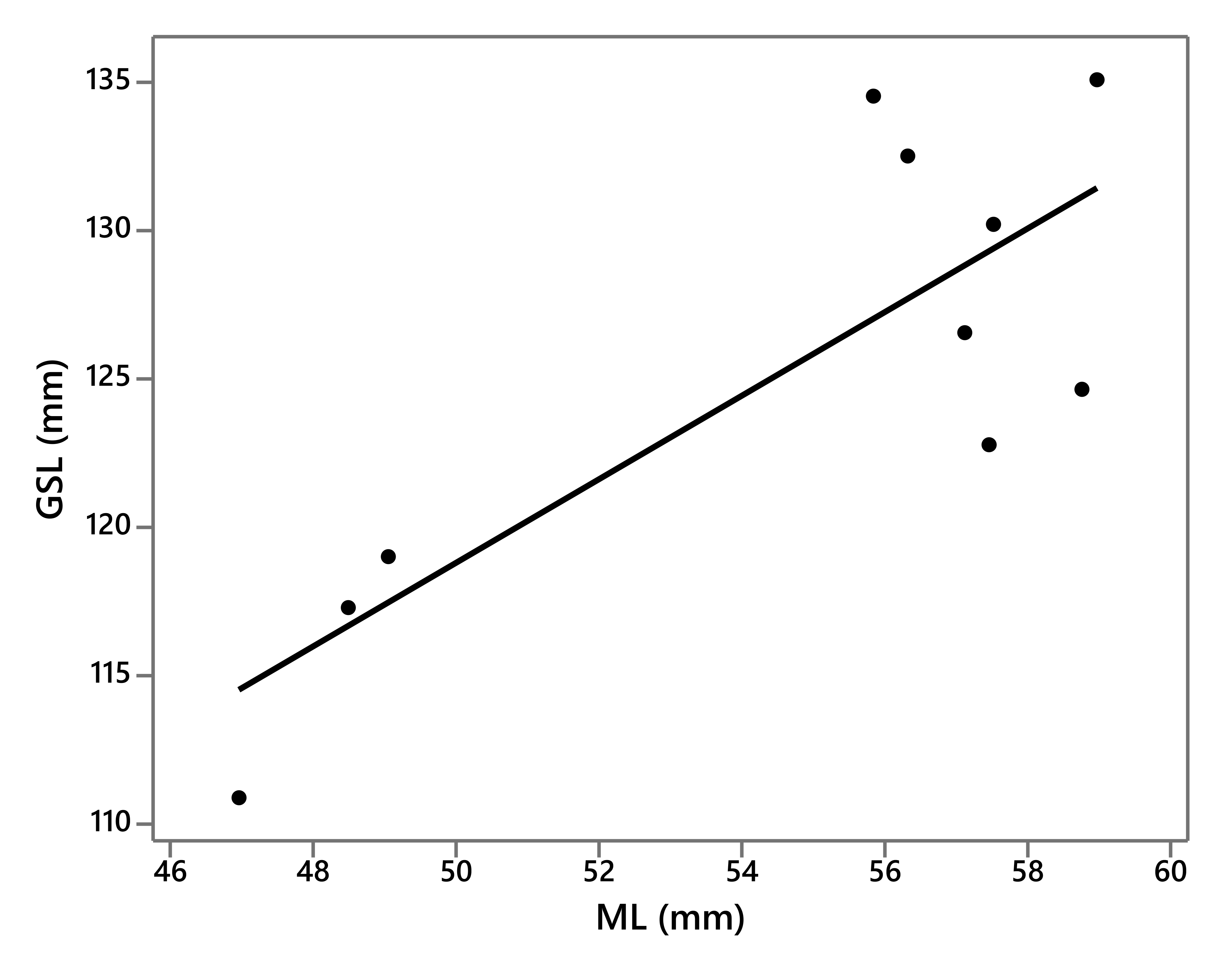

**Fig. 1.** Least-squares regression plot of ML against GSL from the dataset of *Tamandua* in the Table 1.

Correlation coefficient (r) = 0.809

R-squared = 65.457%

P-value of the ANOVA = 0.0046

Equation of the fitted model: GSL = 48.413 + 1.408*ML

There is a strong positive linear correlation between GSL and ML. This relationship is statistically significant at the 95% confidence level.

Estimated GSL for Gen. et sp. nov. (VPPLT 975; ML = 49.87) = 118.623

**Table 2.** Distribution of GSL and body mass values of 10 specimens of *Tamandua tetradactyla* from Suriname, northeastern South America, reported by Husson (1978). The collection numbers belong to the old Museum of Leiden, current Naturalis Biodiversity Center, the Netherlands.

| **Specimen** | **GSL** | **Body mass (g)** |
| --- | --- | --- |
| 22552 | 135.7 | 6000 |
| 10455 | 128.6 | 3750 |
| 17770 | 134.4 | 6500 |
| 17801 | 114.8 | 3750 |
| 18189 | 133.7 | 5500 |
| 18190 | 106 | 2500 |
| 18191 | 129.5 | 7500 |
| 18192 | 127.3 | 5250 |
| 18193 | 126.5 | 5000 |
| 18194 | 133.2 | 5500 |

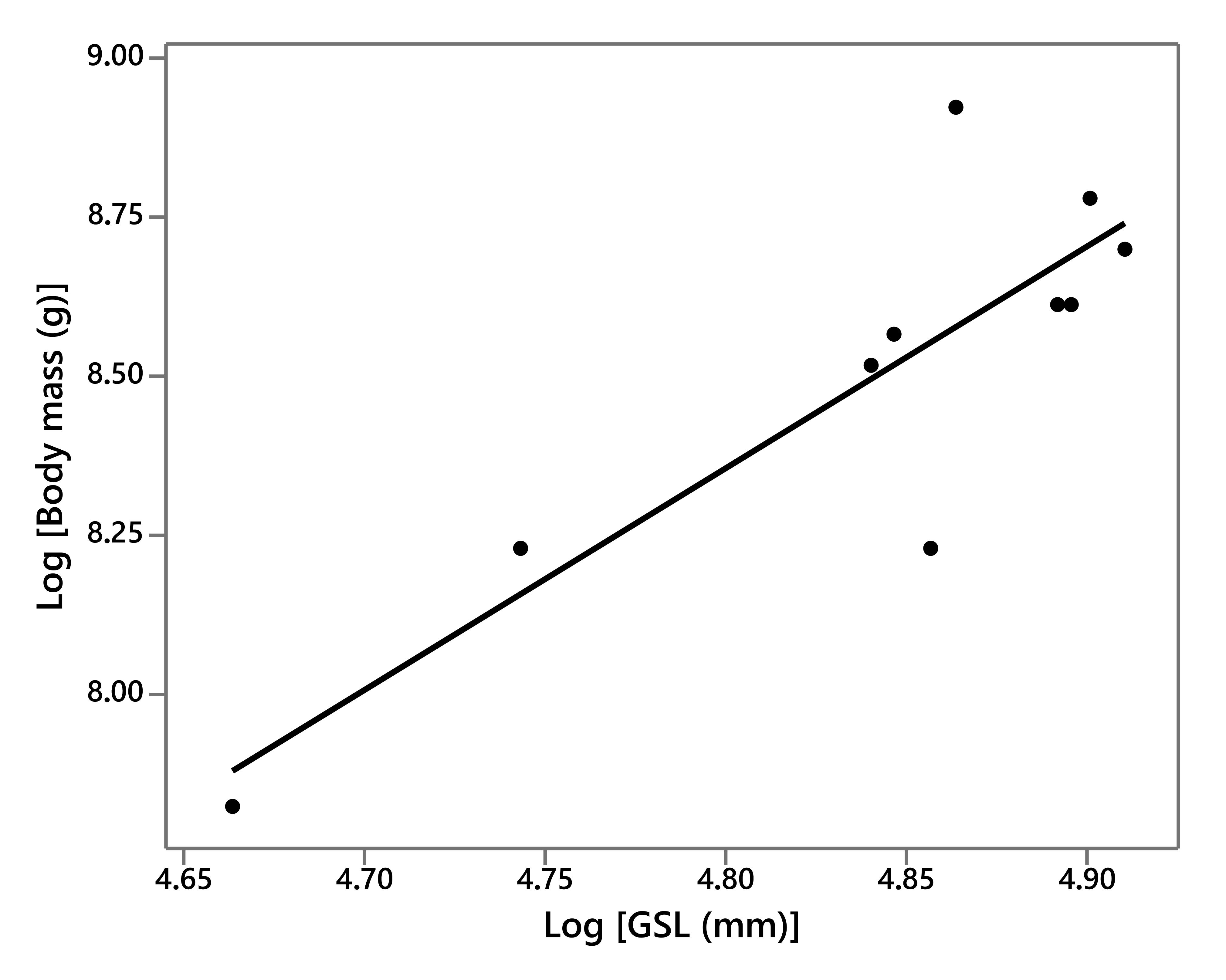

**Fig. 2.** Least-squares regression plot of Log (GSL) against Log (body mass) from the dataset of *Tamandua* in the Table 2.

Correlation coefficient (r) = 0.854

R-squared = 72.929%

P-value of the ANOVA = 0.0017

Equation of the fitted model: Log (body mass) = - 8.369 + 3.484*Log (GSL)

There is a strong positive linear correlation between body mass and GSL. This relationship is statistically significant at the 95% confidence level.

Estimated body mass for Gen. et sp. nov. (VPPLT 975) = 3911.9 g
